# MDA5 ISGylation is crucial for immune signaling to control viral replication and pathogenesis

**DOI:** 10.1101/2024.09.20.614144

**Authors:** Lucky Sarkar, GuanQun Liu, Dhiraj Acharya, Junji Zhu, Zuberwasim Sayyad, Michaela U. Gack

## Abstract

The posttranslational modification (PTM) of innate immune sensor proteins by ubiquitin or ubiquitin-like proteins is crucial for regulating antiviral host responses. The cytoplasmic dsRNA receptor melanoma differentiation-associated protein 5 (MDA5) undergoes several PTMs including ISGylation within its first caspase activation and recruitment domain (CARD), which promotes MDA5 signaling. However, the relevance of MDA5 ISGylation for antiviral immunity in an infected organism has been elusive. Here, we generated knock-in mice (*MDA5^K23R/K43R^*) in which the two major ISGylation sites, K23 and K43, in MDA5 were mutated. Primary cells derived from *MDA5^K23R/K43R^* mice exhibited abrogated endogenous MDA5 ISGylation and an impaired ability of MDA5 to form oligomeric assemblies leading to blunted cytokine responses to MDA5 RNA-agonist stimulation or infection with encephalomyocarditis virus (EMCV) or West Nile virus. Phenocopying *MDA5^−/−^* mice, the *MDA5^K23R/K43R^* mice infected with EMCV displayed increased mortality, elevated viral titers, and an ablated induction of cytokines and chemokines compared to WT mice. Molecular studies identified human HERC5 (and its functional murine homolog HERC6) as the primary E3 ligases responsible for MDA5 ISGylation and activation. Taken together, these findings establish the importance of CARD ISGylation for MDA5-mediated RNA virus restriction, promoting potential avenues for immunomodulatory drug design for antiviral or anti-inflammatory applications.

**Significance Statement:** The work by many groups demonstrated the important role of ubiquitination in modulating the activity of innate immune sensors. In contrast, little is still known about the significance of ISGylation in immune receptor regulation. In this study, we generated knock-in mice in which the two major ISGylation sites of the RNA sensor MDA5 were mutated. Cells from these MDA5-ISGylation-defective mice showed impaired MDA5 oligomerization and antiviral signaling as compared to WT mice. Virus-infected MDA5 knock-in mice displayed ablated antiviral responses, uncontrolled viral replication, and higher mortality. Our study identified HERC5 as the E3 ligase responsible for MDA5 ISGylation and activation. These data may offer opportunities for immune-based antiviral design or ways to alleviate inflammatory diseases associated with overzealous MDA5 activation.

## Introduction

Innate immune surveillance serves as the body’s first line of defense mechanism against a plethora of intruding pathogens whereby pathogen-associated molecular patterns (PAMPs) such as viral RNA and DNA are recognized (1–3). Upon sensing pathogenic ‘non-self’ nucleic acids, germline-encoded pattern-recognition receptors (PRRs) expressed in innate immune (*e.g.,* macrophages) and non-immune (*e.g.,* epithelial or fibroblast) cells confer an amplitude of host antiviral responses. These include *1)* type I or III interferon (IFN)-mediated immunity, *2)* the induction of proinflammatory cytokines, and *3)* upregulation of IFN-stimulated genes (ISGs) in response to type I or III IFN receptor activation and JAK-STAT1/2 signaling. Ultimately, this complex innate immune program initiated by PRRs leads to the activation of adaptive immunity (typically mediated by T and B cells) (4).

Innate immunity in response to viral RNA sensing in the cytoplasm is orchestrated by several receptor proteins, primarily the RIG-I-like receptors (RLRs) retinoic acid-inducible gene-I (RIG-I) and MDA5 (5). These RNA helicases detect specific RNA species, such as 5’-triphosphate-containing RNA (RIG-I) or longer and more complex dsRNA structures (MDA5), after RNA virus infections. Besides RNA viruses, herpesviruses and adenoviruses also activate RLRs where either viral RNAs or certain mislocalized or modified host RNAs harboring signature immunostimulatory features (*i.e.,* 5’-triphosphate moiety and dsRNA portions) are recognized (6, 7). This RNA sensing event then triggers a signaling cascade that is mediated by mitochondrial antiviral-signaling protein (MAVS) and the TBK1-IRF3/7 axis, promoting a transcriptional program comprising IFNs, antiviral effectors (typically the gene products of IFN-stimulated genes (ISGs)), and proinflammatory cytokine or chemokine molecules (5, 8). The antiviral program induced by RLRs ultimately suppresses the replication of diverse RNA viruses (such as flaviviruses, influenza viruses, and coronaviruses) and can also prompt tissue inflammation (9).

Protein posttranslational modifications (PTMs) modulate the physiological functions of cells by altering protein conformation, activity, stability, and/or localization (10, 11). In particular, innate immune sensors are intricately regulated by a ‘PTM-code’ which determines the timing and/or magnitude of PRR activation (5, 12). On the other hand, PTMs can also negatively regulate sensor activation, curbing excessive cytokine responses that can lead to deleterious outcomes such as autoimmune conditions. Serine/Threonine phosphorylation and lysine ubiquitination are the most well-characterized PTMs regulating RLR activity (5). In unstimulated or uninfected cells, MDA5 and RIG-I are phosphorylated in their N-terminal caspase activating and recruitment domains (CARDs) and C-terminal domain (CTD) (13–16). CARD dephosphorylation by a phosphatase complex comprised of protein phosphatase 1 alpha or gamma (PP1α/γ) and the RIG-I/MDA5-targeting subunit PPP1R12C, allows for transition from their signaling-restrained states to signal-transducing ‘active’ forms (14, 17). Specifically, RNA virus infection releases PPP1R12C tethered to actin filaments, allowing its recruitment to RIG-I and MDA5 as part of a catalytically active PP1 complex to dephosphorylate the RLR CARDs. Similarly, the CTD of RLRs is dephosphorylated after RNA virus infection (17). Dephosphorylated RIG-I then undergoes TRIM25- and Riplet-mediated K63-linked polyubiquitination in its CARDs and CTD, respectively (18). These polyubiquitination modifications promote and stabilize RIG-I oligomer formation and thereby its activation to initiate signaling via MAVS (5). MDA5 was shown to undergo K63-linked ubiquitination in its helicase domain catalyzed by the E3 ubiquitin ligase TRIM65, which facilitates MDA5 activation and downstream signaling (19). Whether the MDA5 CARDs undergo K63-linked ubiquitination in cells (vs. cell-free systems) has been controversial (5), prompting research investigations into activating PTMs in the MDA5 CARDs triggered by MDA5 dephosphorylation. Our recent study revealed that MDA5 dephosphorylation induces MDA5 CARD ISGylation (*i.e.,* conjugation with the ubiquitin-like protein ISG15) at two major sites, K23 and K43 (20). MDA5 ISGylation drives antiviral IFN responses restricting a range of RNA viruses including encephalomyocarditis virus (EMCV), Zika virus, and severe acute respiratory syndrome coronavirus 2 (SARS-CoV-2) in human cells (20). Conversely, as a viral tactic evolved to escape ISGylation-dependent MDA5 signaling, the SARS-CoV-2 papain-like protease (PLpro) actively removes ISG15 from the MDA5 CARDs (20, 21). The physiological function of MDA5 ISGylation at the endogenous protein level and its *in vivo* relevance for controlling virus infection, however, have not yet been elucidated.

In this study, we generated *MDA5^K23R/K43R^* knock-in mice and showed that the combined mutation of K23 and K43 ablated endogenous MDA5 ISGylation and oligomerization and thereby MDA5-mediated antiviral cytokine responses, leading to uncontrolled RNA virus-induced pathogenesis. Furthermore, we identified human HERC5 (or HERC6, the functional murine homolog) as the E3 ligase enzyme responsible for catalyzing MDA5 ISGylation, enabling MDA5 activation and antiviral signaling.

## Results

### Ablated MDA5 ISGylation and oligomerization in cells from *MDA5^K23R/K43R^* mice

Our previous work indicated that human MDA5 (hMDA5) undergoes ISGylation at K23 and K43 in the first CARD and that ISGylation promotes MDA5 signaling ability (20). As K23 and K43 are highly conserved in MDA5 across mammalian species including mice (***SI Appendix,* Fig. S1*A***), we sought to determine the physiological relevance of MDA5 CARD ISGylation at the endogenous protein level and for host antiviral defense *in vivo*. To this end, we generated MDA5 knock-in mice (termed *MDA5^K23R/K43R^*) by introducing the K23R and K43R mutations into the native *Mda5*/*Ifih1* locus using CRISPR-Cas9 technology and a targeting repair vector containing the double mutant exon 1 to replace the WT exon 1 (**Fig. 1*A B*, *SI Appendix* Fig. S1*B* and Methods**). In parallel, *MDA5^−/−^* mice in which the exon 1 genomic region was deleted due to non-homologous end joining (NHEJ) were generated as a matched control. All mouse lines were screened and validated using a three-set PCR genotyping strategy and by genomic DNA sequencing (**Fig. 1*B**, SI Appendix* Fig. S1*B* and Methods**).

**Fig. 1.**
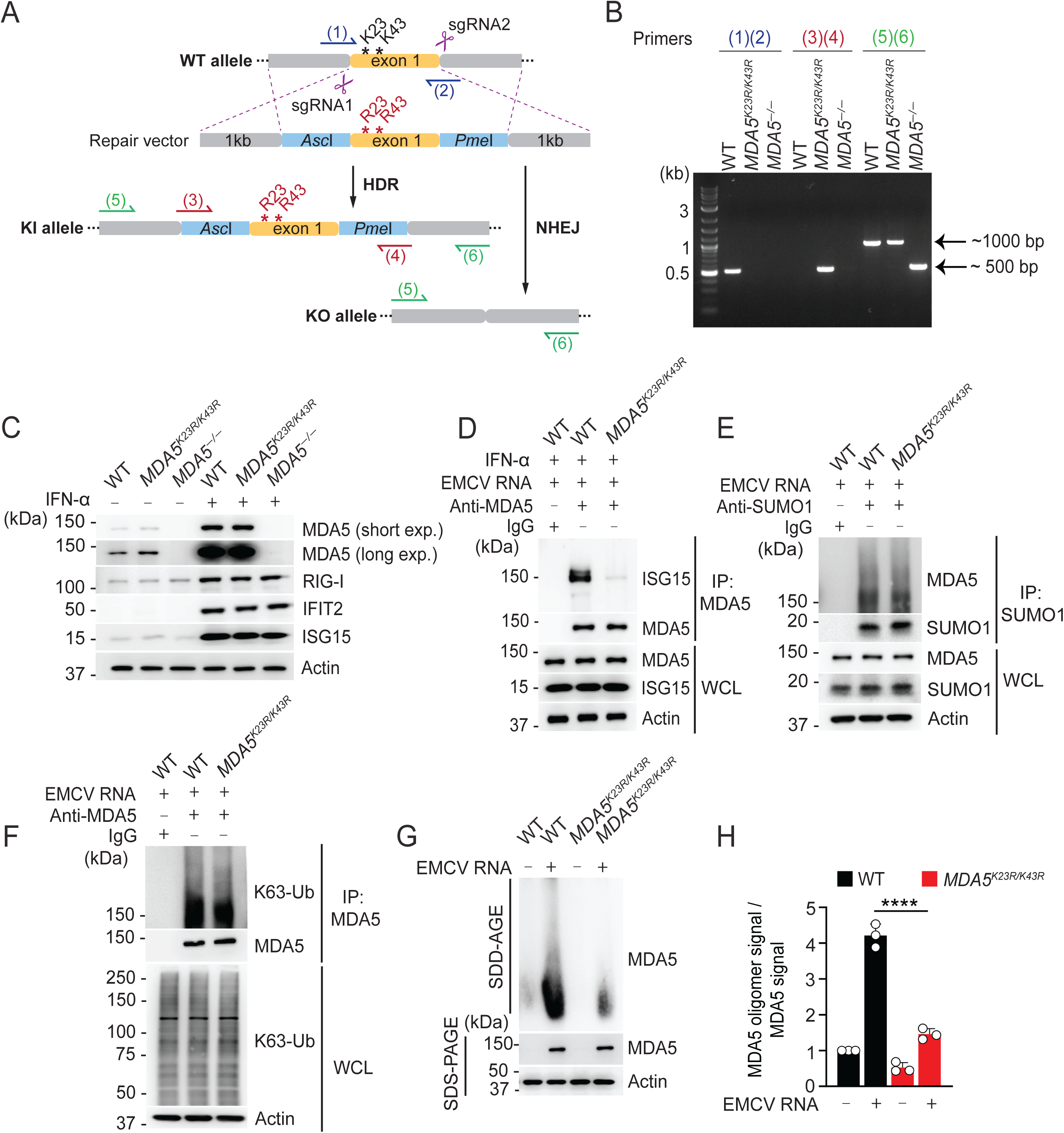
Impaired MDA5 ISGylation and oligomerization in *MDA5^K23R/K43R^* mouse cells. ***(A)*** Schematic of the CRISPR-Cas9 editing strategy for the generation of WT, *MDA5^K23R/K43R^*, and *MDA5^−/−^* mice. See methods and *SI Appendix*, Fig. *S1B* for details. The two conserved lysine residues (K23 and K43; codons AAA and AAA; black asterisks) were mutated to arginines (K23R/K43R; codons AGA and AGA; red asterisks). HDR, homology-directed repair. NHEJ, non-homologous end-joining. sgRNA, single-guide RNA. ***(B)*** Schematic diagram of the validation strategy of the transgenic mouse lines by PCR genotyping. Genomic DNA isolated from ear tissue was amplified to detect the presence of *Mda5/Ifih1* mutant (K23R/K43R) exon1 locus using the indicated primers by agarose gel electrophoresis. The primer pair (1) and (2) generates a 523 bp-fragment in WT mice; the primer pair (3) and (4) generates a 537 bp-fragment in *MDA5^K23R/K43R^* mice; and the primer pair (5) and (6) generates a 1046 bp-fragment in WT mice, a 1062 bp-fragment in *MDA5^K23R/K43R^* mice, and a 565 bp-fragment in *MDA5^−/−^* mice. ***(C)*** Analysis of the protein abundance of endogenous MDA5, RIG-I, and downstream ISGs (IFIT2 and ISG15) in the whole cell lysates (WCLs) of primary mouse dermal fibroblasts (MDFs) isolated from 6-8-week-old WT, *MDA5^K23R/K43R^*, and *MDA5^−/−^* mice that were stimulated *ex vivo* with IFN-α (500 U/mL) for 24 h or that remained untreated (−), determined by immunoblot (IB) analysis. ***(D)*** Endogenous MDA5 ISGylation in MDFs from WT or *MDA5^K23R/K43R^* mice that were pre-stimulated for 8 h with IFN-α (1000 U/mL) and then transfected with EMCV RNA (0.4 µg/mL) for 16 h to stimulate MDA5 activation, determined by IP with anti-MDA5 (or an IgG isotype control) and IB with anti-ISG15. ***(E)*** Endogenous MDA5 SUMOylation in WT or *MDA5^K23R/K43R^* mouse-derived MDFs transfected with EMCV-RNA (0.4 µg/mL) for 16 h, determined by IP with anti-SUMO1 (or an IgG isotype control) and IB with anti-MDA5. ***(F)*** K63-linked ubiquitination of endogenous MDA5 in MDFs from WT or *MDA5^K23R/K43R^* mice that were transfected with EMCV RNA (0.4 µg/mL) for 16 h, determined by IP with anti-MDA5 and IB with K63-polyubiquitin-linkage-specific antibody (K63-Ub). ***(G)*** Endogenous MDA5 oligomerization in WT and *MDA5^K23R/K43R^* mouse-derived MDFs that were transfected with EMCV RNA (0.4 μg/mL) for 8 h, assessed by SDD-AGE and IB with anti-MDA5. Equal protein abundance of MDA5 in WT and *MDA5^K23R/K43R^* mouse cells was validated by SDS-PAGE and IB with anti-MDA5 (with Actin as loading control). ***(H)*** Densitometric analysis of the MDA5 oligomer signal, normalized to the respective MDA5 protein abundance, for the experiment in (G). Values represent relative signal intensity normalized to values for unstimulated WT control cells, set to 1. Data are representative of at least two independent experiments (mean ± s.d. of n = 3 biological replicates in [H]). ****P < 0.0001 (two-tailed, unpaired student’s *t*-test).

We next assessed the protein abundance of endogenous MDA5 in primary mouse dermal fibroblasts (MDFs) isolated from the three mouse lines both in unstimulated (basal) conditions and after exogenous IFN-α stimulation (**Fig. 1*C***). This showed comparable endogenous MDA5 protein expression in the cells from WT and *MDA5^K23R/K43R^* mice, and further, confirmed the absence of MDA5 expression in the cells from *MDA5^−/−^* mice. Notably, equal RIG-I and downstream ISG (*i.e.,* IFIT2 and ISG15) protein expression was observed after IFN-α stimulation in the MDFs from all three mouse lines (**Fig. 1*C***), demonstrating intact IFN-α/β receptor (IFNAR) signaling. Next, we tested the ISGylation of endogenous MDA5 after stimulation with EMCV RNA, a specific agonist of MDA5 (5, 22), in MDFs isolated from *MDA5^K23R/K43R^* mice and WT mice (**Fig. 1*D***). Of note, experimental conditions were used where ISG15 protein expression was comparable in both WT and knock-in mouse cells, allowing us to unambiguously compare the ISGylation of WT and mutant MDA5. Cells from WT mice showed robust endogenous MDA5 ISGylation after EMCV RNA stimulation. In contrast, EMCV RNA-stimulated cells from *MDA5^K23R/K43R^* mice exhibited a near-abolished ISGylation of endogenous MDA5 (**Fig. 1*D***). Importantly, the levels of K63-linked polyubiquitination and SUMOylation of endogenous MDA5 (19, 23) in cells from *MDA5^K23R/K43R^* and WT mice were comparable (**Fig. 1***E −F*), strengthening our previous data (20) that showed that the mutation of K23 and K43 specifically abrogates ISG15 conjugation but does not affect —directly or indirectly— MDA5 ubiquitination or SUMOylation.

Upon binding to dsRNA in the cytosol, hMDA5 is primed by CARD ISGylation facilitating its multimerization (20). Consistent with these previous findings on exogenous WT and K23R/K43R hMDA5, endogenous mMDA5 exhibited efficient oligomerization in EMCV RNA-stimulated MDFs from WT control mice; however, endogenous mMDA5 oligomerization was substantially impaired in cells derived from *MDA5^K23R/K43R^* mice (**Fig. 1*G H*, and *SI Appendix*, Fig. S1*C−D).*** Collectively, these findings show that endogenous MDA5 undergoes ISGylation at K23 and K43, which is important for its ability to oligomerize in response to RNA agonist stimulation.

### MDA5 ISGylation is pivotal for eliciting IFN and ISG responses against picornavirus infection in fibroblasts

To elucidate the role of MDA5 ISGylation in downstream signal transduction, we assessed specific activating phosphorylation marks for STAT1 (downstream of IFNAR) as well as IRF3 and TBK1 (both downstream of MDA5 and other PRRs) in MDF cells derived from WT and *MDA5^K23R/K43R^* mice upon infection with EMCV. Cells from *MDA5^−/−^* mice were included as a control. EMCV-infected cells from WT mice, but not *MDA5^K23R/K43R^* and *MDA5^−/−^* mice, exhibited robust STAT1 phosphorylation (**Fig. 2*A***). In accord, TBK1 and IRF3 phosphorylation was effectively elicited in cells from WT mice following EMCV infection. In contrast, cells derived from *MDA5^K23R/K43R^* and *MDA5^−/−^* mice showed impaired activating phosphorylations for TBK1 and IRF3 (***SI Appendix*, Fig. S2*A***). Importantly, MDFs from WT, *MDA5^K23R/K43R,^* and *MDA5^−/−^* mice showed comparable TBK1, IRF3, and STAT1 phosphorylations upon infection with Sendai virus (SeV, a virus that is sensed by RIG-I), demonstrating the integrity of the RIG-I signaling pathway in the cells derived from *MDA5^K23R/K43R^* and *MDA5^−/−^* mice. Consistent with these data, the transcript expression of type I IFN (*i.e., Ifna1),* ISGs (*i.e., Mx1* and *Oas1b*), and proinflammatory cytokines and chemokines (*i.e., Tnf, Ccl5, and Cxcl10*) were efficiently elicited in MDFs from WT mice over a time course of EMCV RNA stimulation. In comparison, antiviral and proinflammatory gene induction was impaired in EMCV RNA-transfected cells from *MDA5^K23R/K43R^* and *MDA5^−/−^* mice. Notably, *MDA5^−/−^* mouse cells consistently showed a stronger diminishment of antiviral gene induction compared with the cells from *MDA5^K23R/K43R^* mice (**Fig. 2*B−D* and *SI Appendix*, Fig. S2*B−D***). MDFs derived from WT, *MDA5^K23R/K43R,^* and *MDA5^−/−^* mice, however, responded equally well to rabies virus leader RNA (RABV_Le_; an RNA agonist activating RIG-I (24)) (**Fig. 2*B−D* and *SI Appendix,* Fig. S2*B−D***). Consistent with these data using RLR RNA-ligands, authentic EMCV infection in cells from WT mice, but not in cells from *MDA5^K23R/K43R^* and *MDA5^−/−^* mice, effectively elicited antiviral gene responses, while SeV infection robustly stimulated an antiviral response in the cells from all three mouse lines (***SI Appendix,* Fig. S2*E−I***). Consistent with our data on antiviral gene induction, we observed strongly diminished and ablated IFN-β protein secretion in MDFs derived from *MDA5^K23R/K43R^* and *MDA5^−/−^* mice, respectively (compared to cells from WT mice) after MDA5, but not RIG-I, stimulation (**Fig. 2*E−F***). These results indicate that ISGylation of endogenous MDA5 is required for its functional ability to instigate an antiviral cellular defense program.

**Fig. 2.**
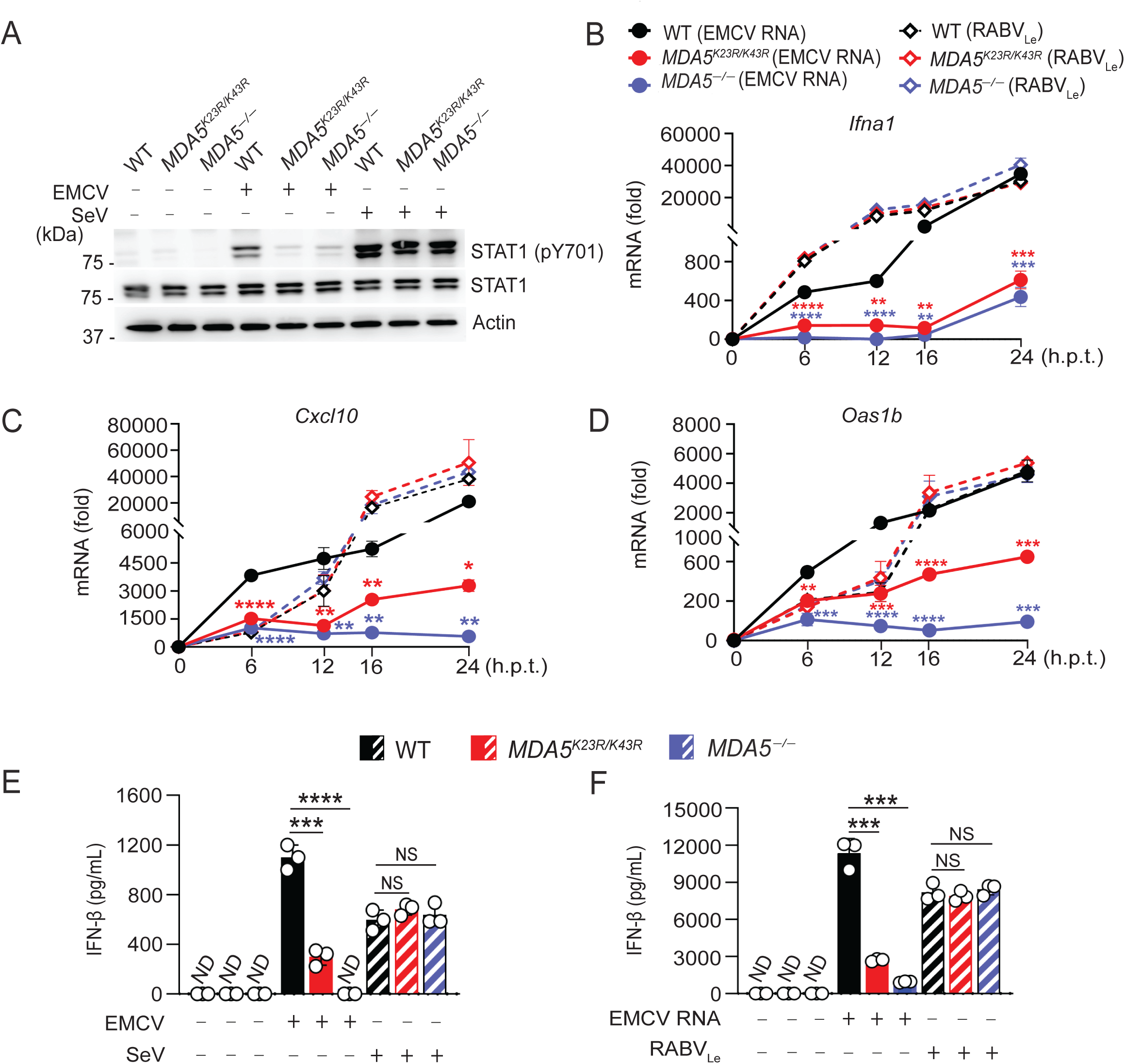
Ablated MDA5 antiviral signaling in *MDA5^K23R/K43R^* mouse-derived dermal fibroblasts. ***(A)*** STAT1 phosphorylation in WT, *MDA5^K23R/K43R^*, and *MDA5^−/−^* mouse-derived MDFs that were infected for 12 h with either EMCV (MOI 2) or SeV (250 hemagglutination units [HAU]/mL) or that remained uninfected (−), assessed in the WCLs by IB with anti-pT701-STAT1 and anti-STAT1. ***(B−D)*** *Ifna1*, *Cxcl10,* and *Oas1b* transcript abundance in WT, *MDA5^K23R/K43R^*, and *MDA5^−/−^* mouse-derived MDFs that were transfected with EMCV RNA (0.4 μg/mL) or RABV_Le_ (1 pmol/mL) for the indicated times, determined by RT-qPCR. ***(E−F)*** Secreted IFN-β protein in the supernatant of WT*, MDA5^K23R/K43R^*, and *MDA5^−/−^* mouse-derived MDFs that were either mock-treated (−) (E and F), infected for 12 h with EMCV (MOI 1) or SeV (20 HAU/mL) (E), or transfected for 12 h with EMCV RNA (0.4 µg/mL) or RABV_Le_ (1 pmol/mL) (F), determined by ELISA. Data are representative of at least two independent experiments (mean ± s.d. of n = 3 biological replicates in B−F). *P < 0.05, **P < 0.01, ***P < 0.001, and ****P < 0.0001 (two-tailed, unpaired student’s *t*-test). Red and blue asterisks in (B−D) indicate the statistical significance (P-values) for WT vs. *MDA5^K23R/K43R^* and WT vs. *MDA5^−/−^* samples, respectively. h.p.t., hours post-transfection. ND, not detected. NS, statistically not significant.

### ISG15 conjugation of the MDA5 CARDs is required for innate signaling in immune cells

We next sought to determine the role of CARD ISGylation in MDA5 signaling in immune cells, in particular primary bone marrow-derived macrophages (BMDMs). Similar to our results obtained from MDFs, EMCV-infected BMDMs from *MDA5^K23R/K43R^* and *MDA5^−/−^* mice exhibited strongly diminished phosphorylation of IRF3, TBK1, and STAT1 compared to BMDMs from WT mice (**Fig. 3*A* and *SI Appendix,* Fig. S3*A***). In accord, cytokine and chemokine gene expression upon EMCV infection or EMCV-RNA transfection was impaired in *MDA5^K23R/K43R^* cells compared to WT control cells (**Fig. 3*B−D* and *SI Appendix*, Fig. S3*B−C***). In stark contrast, the signaling molecule activation and antiviral gene responses of SeV-infected or RABV_Le_-transfected *MDA5^K23R/K43R^* mouse-derived BMDMs were comparable to those in cells from WT mice (**Fig. 3*B−D* and *SI Appendix*, Fig. S3*B−C***). These results show that immune cells derived from *MDA5^K23R/K43R^* mice exhibit abrogated MDA5 antiviral signaling.

**Fig. 3.**
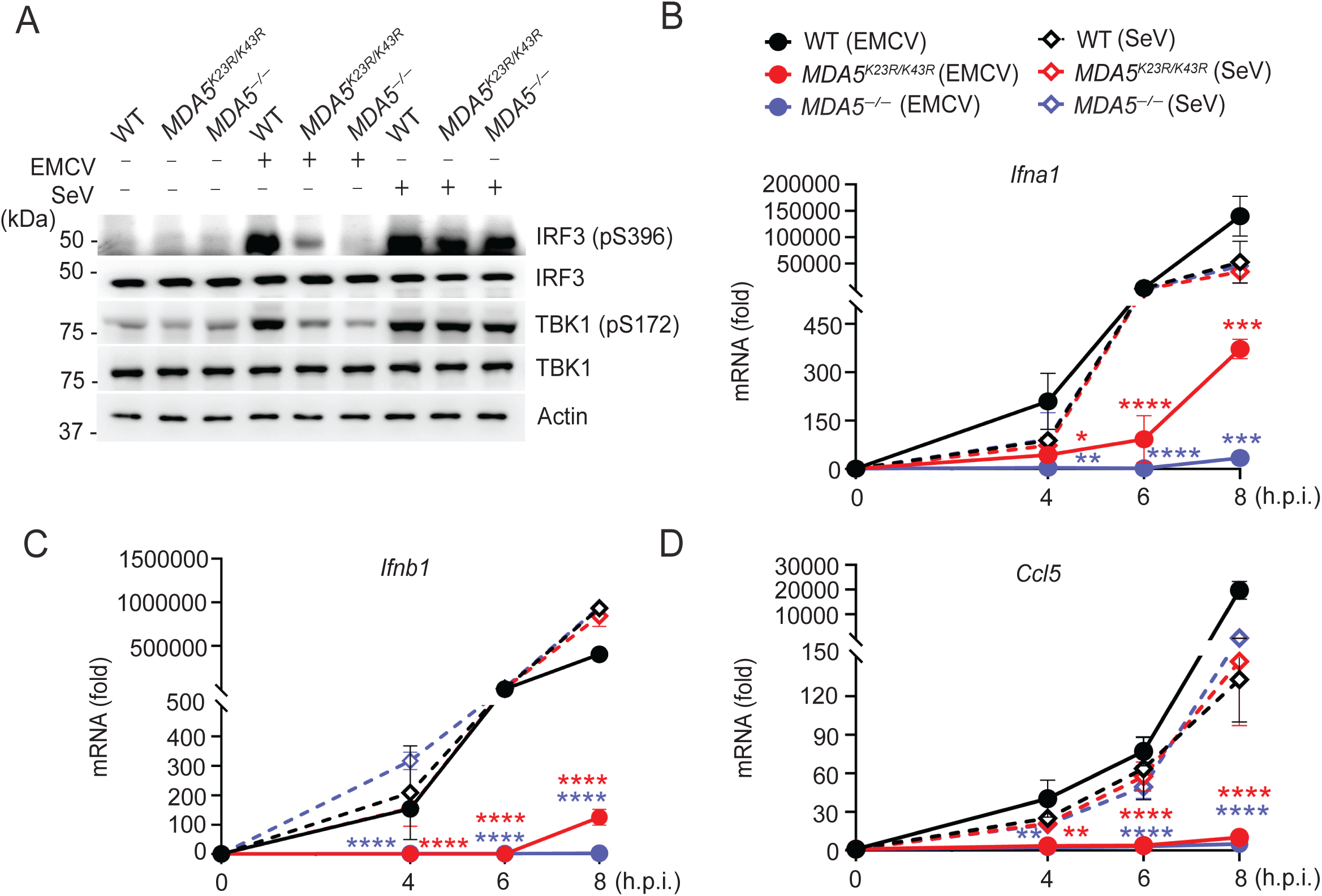
MDA5 signaling to EMCV, but not SeV, infection is impaired in *MDA5^K23R/K43R^* mouse-derived immune cells. ***(A)*** Phosphorylation of endogenous IRF3 and TBK1 in WT*, MDA5^K23R/K43R^*, and *MDA5^−/−^* mouse-derived BMDMs that were infected for 6 h with either EMCV (MOI 5) or SeV (200 HAU/mL), assessed in the WCLs by IB with anti-pS396-IRF3 and anti-pS172-TBK1. WCLs were further immunoblotted with anti-IRF3, anti-TBK1, and anti-Actin (loading control). ***(B−D)*** *Ifna1, Ifnb1,* and *Ccl5* transcripts in WT, *MDA5^K23R/K43R^*, and *MDA5^−/−^* mouse-derived BMDMs that were infected with either EMCV (MOI 1) or SeV (20 HAU/mL) for the indicated times. Data are representative of at least two independent experiments (mean ± s.d. of n = 3 biological replicates in B−D). *P < 0.05, **P < 0.01, ***P < 0.001, and ****P < 0.0001 (two-tailed, unpaired student’s *t*-test). Red and blue asterisks in (B−D) indicate the statistical significance (P-values) for WT vs. *MDA5^K23R/K43R^* and WT vs. *MDA5^−/−^* samples, respectively. h.p.i., hours post-infection.

### MDA5 ISGylation is important for eliciting an antiviral transcriptional program against coronaviruses and flaviviruses

In addition to detecting picornavirus infections, MDA5 is a major receptor for sensing coronaviruses and flaviviruses. As such, we investigated the requirement of MDA5 ISGylation at K23 and K43 for initiating an innate transcriptional program to stimulation with SARS-CoV-2 (coronavirus) RNA and to authentic West Nile virus (WNV, a flavivirus) infection. Transfection of SARS-CoV-2 RNA (which activates primarily MDA5 (20)) into MDFs from *MDA5^K23R/K43R^* mice and *MDA5^−/−^* mice, respectively, severely impaired and abrogated, antiviral and proinflammatory gene expression as compared to that induced in WT cells (**Fig. 4*A***). Moreover, *MDA5^K23R/K43R^* or *MDA5^−/−^* mouse-derived MDFs exhibited blunted antiviral transcriptional responses following WNV infection as compared to control cells (**Fig. 4*B***). Of note, in these experiments, we measured antiviral gene induction specifically at a late time (*i.e.,* 60 h) in WNV infection where MDA5 was shown to play a major role in flaviviral RNA detection, whereas RIG-I senses WNV early in infection (25). Together with our data on EMCV, these findings strengthen the importance of CARD ISGylation for MDA5’s ability to elicit an innate immune program against RNA viruses from diverse families.

**Fig. 4.**
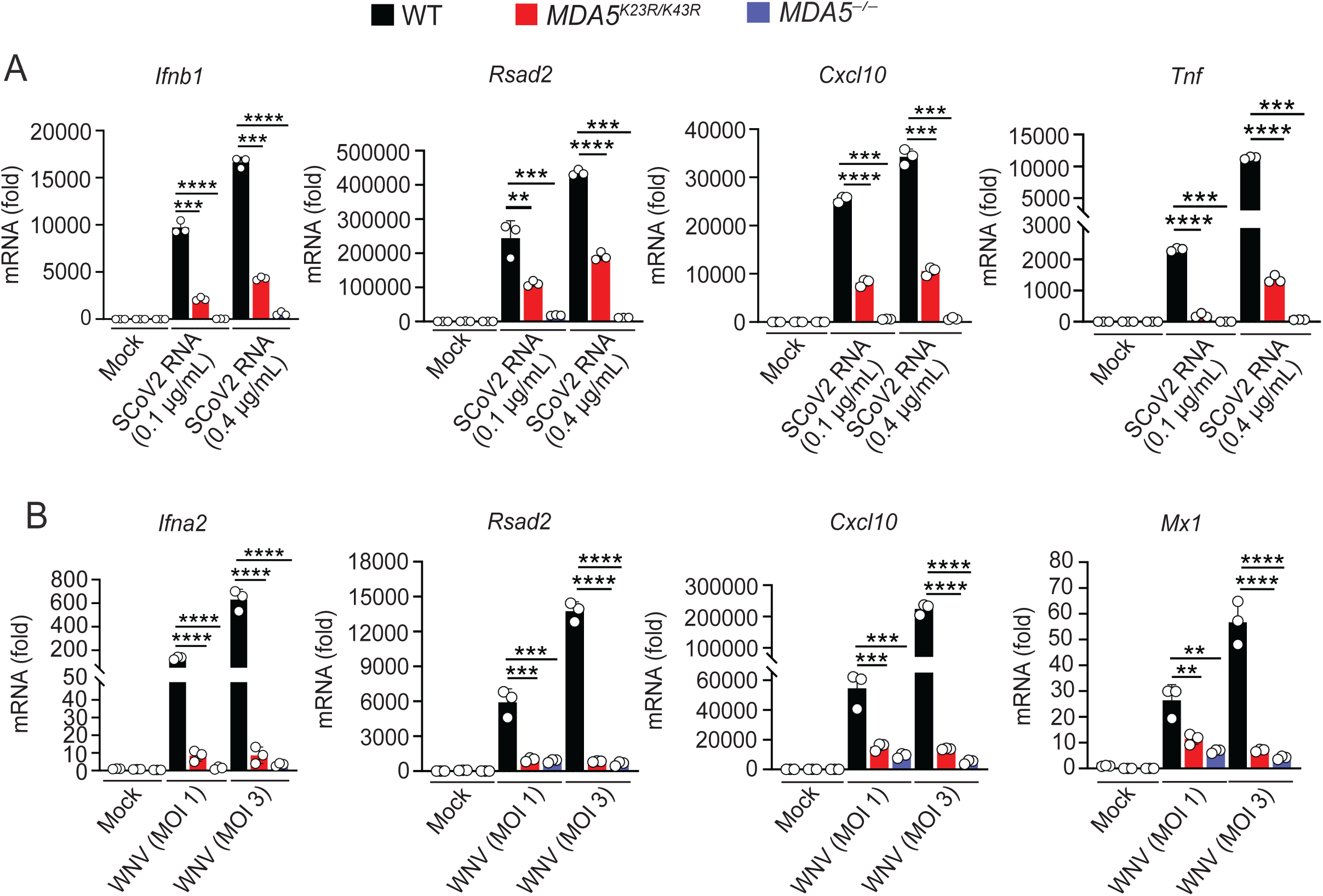
*MDA5^K23R/K43R^* mouse-derived cells are deficient in mounting an innate immune response to coronavirus or flavivirus challenge. ***(A)*** RT-qPCR analysis of the indicated antiviral or proinflammatory gene transcripts in WT, *MDA5^K23R/K43R^*, and *MDA5^−/−^* mouse-derived MDFs at 16 h post-transfection with SARS-CoV-2 RNA (0.1 or 0.4 µg/mL). Mock-treated cells served as control. ***(B)*** RT-qPCR analysis of the indicated genes in WT, *MDA5^K23R/K43R^*, and *MDA5^−/−^* mice-derived MDFs that were either mock-treated or infected for 60 h with WNV (MOI 1 or 3). Data are representative of at least two independent experiments (mean ± s.d. of n = 3 biological replicates). **P < 0.01, ***P < 0.001, and ****P < 0.0001 (two-tailed, unpaired student’s *t*-test). SCoV2, SARS-CoV-2.

### HERC5/HERC6 catalyzes MDA5 ISGylation, promoting MDA5 oligomerization and immune signal transduction

To identify the E3 ligase(s) responsible for MDA5 CARD ISGylation, we adopted a candidate approach in which we silenced specific enzymes known to have E3 ligase activity for ISG15 (*i.e.,* HERC5 (26, 27), ARIH1 (Ariadne RBR E3 ubiquitin protein ligase 1 (28)), and TRIM25 (also named estrogen finger protein (EFP) (29)), and tested the effect of silencing on endogenous hMDA5 ISGylation. Knockdown of TRIM65, which mediates the K63-liked ubiquitination of MDA5’s helicase domain (19) and is not known to confer ISG15 E3 ligase activity, served as a control in this experiment. Depletion of endogenous HERC5 ablated MDA5 ISGylation in primary normal human lung fibroblasts (NHLF) as compared to transfection of non-targeting control siRNA (si.C), whereas knockdown of the other E3 ligases had no diminishing effect on MDA5 ISGylation (**Fig. 5*A***). Depletion of endogenous HERC6 (the functional substitute of HERC5 in mice (30, 31)) in primary MDFs near-abolished MDA5 ISGylation induced by EMCV RNA stimulation, to a similar extent as did E1 or E2 silencing (**Fig. 5*B***). In contrast, depletion of endogenous TRIM65 in MDFs did not affect MDA5 ISGylation, ruling out that TRIM65 —either directly or indirectly (for example, via a possible crosstalk between MDA5 K63-linked ubiquitination and ISGylation)— influences MDA5 ISGylation (**Fig. 5*B***). In line with these findings, HERC6 knockdown in EMCV RNA-stimulated WT MDFs noticeably diminished MDA5 oligomerization. By contrast, HERC6 silencing in cells from *MDA5^K23R/K43R^* mice, which showed impaired MDA5 oligomerization (as compared to cells from WT mice), did not further reduce MDA5 oligomerization (**Fig. 5*C* and *D***).

**Fig. 5.**
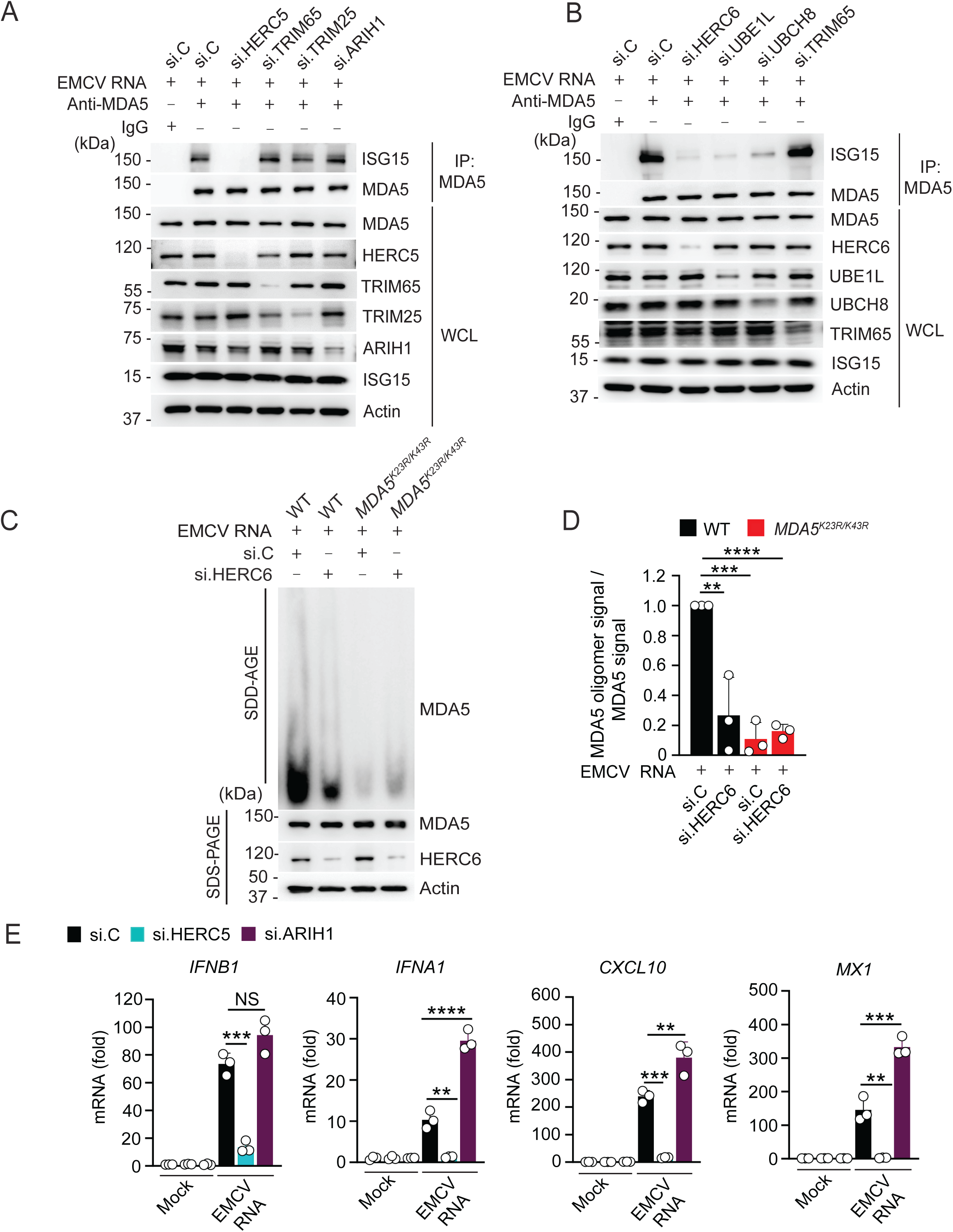
HERC5/HERC6 catalyzes MDA5 ISGylation promoting MDA5 oligomerization and immune signaling. ***(A)*** ISGylation of endogenous MDA5 in primary NHLF cells that were transfected for 48 h with the indicated siRNAs and then transfected with EMCV RNA (0.4 μg/mL) for 16 h, determined by IP with anti-MDA5 (or an IgG isotype control) and IB with anti-ISG15. Knockdown of the individual genes was confirmed in the WCLs by IB with the indicated antibodies. ***(B)*** Endogenous MDA5 ISGylation in WT mouse-derived MDFs that were transfected for 48 h with the indicated siRNAs and then transfected with EMCV RNA (0.4 μg/mL) for 16 h, determined as in (A). Knockdown of the individual genes was confirmed in the WCLs by IB with the indicated antibodies. ***(C)*** Endogenous MDA5 oligomerization in WT and *MDA5^K23R/K43R^* mouse-derived MDFs that were transfected for 48 h with the indicated siRNAs and then transfected with EMCV RNA (0.4 μg/mL) for 16 h, assessed by SDD-AGE and IB with anti-MDA5. Input amounts for MDA5 as well as knockdown of endogenous HERC6 were confirmed by SDS-PAGE and IB with anti-MDA5 or anti-HERC6. ***(D)*** Densitometric analysis of the MDA5 oligomer signal, normalized to the respective MDA5 protein abundance, from the experiment in (C). Values represent relative signal intensity normalized to values for si.C-transfected WT cells, set to 1. ***(E)*** *IFNB1, IFNA1, CXCL10,* and *MX1* gene transcripts in primary NHLF cells that were transfected with the indicated siRNAs and then either Mock-treated or stimulated with EMCV RNA as in (A), determined by RT-qPCR. Data are representative of at least two (A, B, and E) or three (C and D) independent experiments (mean ± s.d. of n = 3 biological replicates in (D and E). *P < 0.05, **P < 0.01, ***P < 0.001, and ****P < 0.0001 (two-tailed, unpaired student’s *t*-test). si.C, non-targeting control siRNA.

Knockdown of HERC5, but not ARIH1, in primary NHLFs markedly reduced the transcript expression of ISGs, cytokines, and chemokines upon EMCV RNA stimulation (**Fig. 5*E* *and SI Appendix, Fig.* S*4A***). Similarly, the knockdown of endogenous HERC6 in WT MDFs abrogated EMCV RNA-induced antiviral gene expression as compared to si.C transfection (***SI Appendix,* Fig. S4*B***). Collectively, these results establish that HERC5 (human) and HERC6 (mouse) are the major E3 ligases that mediate MDA5 ISGylation, ultimately promoting MDA5 oligomerization and antiviral signaling.

### *MDA5^K23R/K43R^* mice are impaired in restricting virus infection

To evaluate the *in vivo* relevance of ISGylation-dependent MDA5 activation in antiviral immunity, we infected WT and *MDA5^K23R/K43R^* mice intraperitoneally with EMCV and monitored morbidity and survival, innate immune responses, and viral titers (**Fig. 6*A***). *MDA5^−/−^* mice were included in these experiments for comparison. *MDA5^K23R/K43R^* and *MDA5^−/−^* mice infected with EMCV exhibited greater body weight loss and accelerated lethality as compared to infected WT mice (**Fig. 6*B* and *SI Appendix,* Fig. S5*A***). Analysis of EMCV replication revealed that *MDA5^K23R/K43R^* and *MDA5^−/−^* mice had significantly higher viral titers in cardiac and brain tissues as compared to WT mice (**Fig. 6*C−D***), indicating enhanced viral replication due to ablated MDA5 activity in the *MDA5^K23R/K43R^* and *MDA5^−/−^* mice. Furthermore, effective IFN-β production was triggered in the blood and heart of infected WT mice. In contrast, IFN-β protein amounts in these tissues were undetectable in infected *MDA5^K23R/K43R^* and *MDA5^−/−^* mice (**Fig. 6*E***). In line with these results, RT-qPCR analysis detected higher viral RNA amounts and strongly reduced cytokine/chemokine transcript levels in the blood (**Fig. 6*F***) and heart (**Fig. 6*G***) of infected *MDA5^K23R/K43R^* mice compared with infected WT control mice. Of note, the impaired antiviral transcriptional program observed for *MDA5^K23R/K43R^* mice was comparable to that of infected *MDA5^−/−^* mice, which also showed blunted cytokine/chemokine induction as expected (**Fig. 6*F−G***). Cumulatively, these results indicate that CARD ISGylation is a key activation mechanism for MDA5 to control RNA virus infection and viral pathogenesis *in vivo*.

**Fig. 6.**
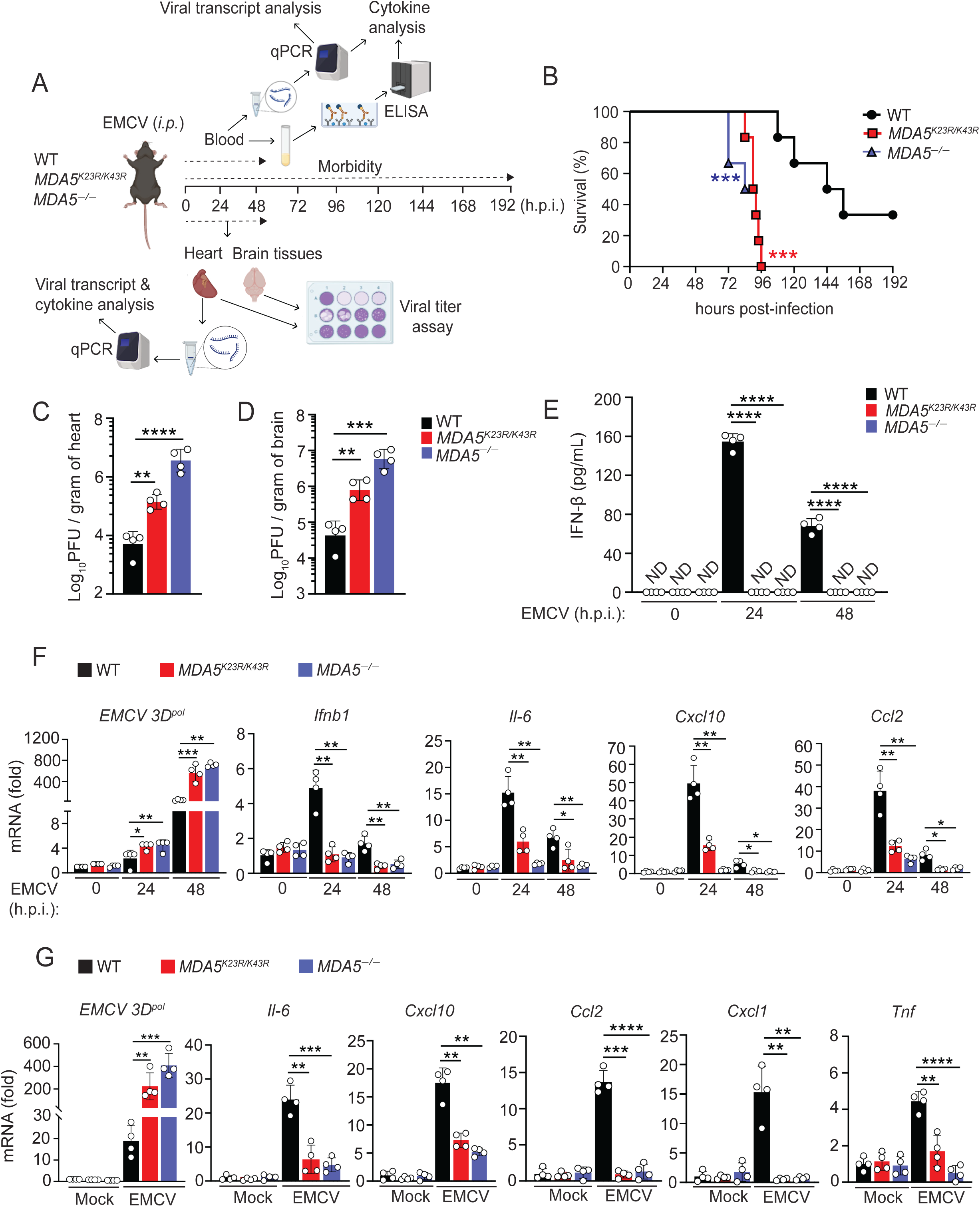
ISGylation-defective *MDA5^K23R/K43R^* mice are impaired in controlling EMCV infection and EMCV-induced pathogenesis. ***(A)*** Overview of the mouse infection studies with EMCV to measure morbidity and survival, viral replication, and cytokine responses. ***(B)*** WT, *MDA5^K23R/K43R^*, and *MDA5^−/−^* mice (6-8-week-old) were infected via intraperitoneal (*i.p.*) inoculation with EMCV (25 PFU). Kaplan-Meier survival curves of EMCV-infected WT, *MDA5^K23R/K43R^*, and *MDA5*^−/−^ mice (n = 6 per genotype). ***(C−G)*** WT, *MDA5^K23R/K43R^*, and *MDA5^−/−^* mice (6-8-week-old) were infected via *i.p.* inoculation with EMCV (10^3 PFU). Viral titers in the heart ***(C)*** and brain ***(D)*** were determined by plaque assay at 48 h p.i., and ***(E)*** IFN-β protein in the blood was analyzed by ELISA at 24 and 48 h.p.i. Furthermore, *EMCV 3D-pol* as well as host antiviral or proinflammatory gene transcripts were measured in blood at 24 and 48 h.p.i. ***(F)*** and in heart tissue at 48 h.p.i. ***(G)***. Data are representative of at least two independent experiments (mean ± s.d. of n = 6 (B) or n = 4 (C−G) biological replicates). *P < 0.05, **P < 0.01, ***P < 0.001, ****P < 0.0001. Mantel-Cox test (B) or two-tailed, unpaired student’s *t*-test (C−G). h.p.i., hours post-infection. ND, non-detected. Parts of Fig. 6A were created using *Biorender.com*.

## Discussion

Fine-tuning the signaling activity of the innate RNA sensor MDA5 has been shown to require several PTMs including phosphorylation, ubiquitination, SUMOylation, and lately, ISGylation (5). While the molecular discoveries on PTM-mediated MDA5 regulation have greatly advanced our understanding of MDA5 activation, the physiological relevance of several of these PTM marks, particularly in an organism, has been elusive. In the present study, we generated *MDA5^K23R/K43R^* mice with mutation of the two key ISGylation sites in MDA5 and investigated the direct contribution of ISGylation for MDA5-dependent antiviral innate immunity. We showed that, like human MDA5, endogenous mouse MDA5 undergoes robust ISGylation, and further, that this modification is crucial for MDA5’s ability to form higher-order oligomeric assemblies and to induce antiviral IFN responses. Notably, this important role of MDA5 CARD ISGylation was observed for various MDA5 stimuli including MDA5-specific RNA ligands (*i.e.,* EMCV-RNA and SARS-CoV-2 RNA) and viruses from different families (*i.e., Picornaviridae* (EMCV) and *Flaviviridae* (WNV), both known to be detected by MDA5). Furthermore, similar to *MDA5^−/−^* mice, *MDA5^K23R/K43R^* mice were highly susceptible to EMCV infection and displayed heightened pathology and lethality owing to diminished antiviral IFN and cytokine/chemokine responses. Our data thus establish ISGylation as a physiologically important PTM governing MDA5 activation and its downstream antiviral signaling.

Our work also identified the E3 ligases catalyzing the CARD ISGylation marks of MDA5. Through a targeted siRNA-based mini-screen, we found that HERC5 and its functional murine homolog, HERC6, represent the key E3 ligases responsible for MDA5 ISGylation, prompting MDA5 downstream antiviral signaling. Interestingly, ISGylation has recently been shown to play important roles in the activation of the cGAS-mediated innate DNA sensing pathway (32–35). HERC5 and mouse HERC6 were also identified to be the critical E3 enzymes involved in the ISGylation of the DNA sensor cGAS and its signaling adaptor STING, promoting HSV-1 restriction (34, 35). These findings highlight HERC5/HERC6-mediated ISGylation as an essential regulatory arm of PRR-induced antiviral innate immunity against both RNA viruses and DNA viruses. While we have not tested directly the *in vivo* role of HERC6 in antiviral defense against MDA5-sensed viruses, a previous study showed that compared to WT mice, *HERC6^−/−^* mice, despite exhibiting ablated global ISGylation, mounted comparable IFN and proinflammatory cytokine responses to infections with SeV and vesicular stomatitis virus, both are known to be primarily sensed by RIG-I. This is consistent with our and others’ observation that ISGylation positively regulates MDA5 signaling but has minimal or even opposing effects on RIG-I activation (20, 36, 37). Future studies are necessary to comprehensively assess the antiviral responses to MDA5- or RIG-I-sensed viruses in *HERC6^−/−^* mice.

Our data strengthened the concept that HERC5/HERC6-mediated ISGylation of the N-terminal CARDs is important for efficient MDA5 oligomerization. Our observation that *MDA5^K23R/K43R^* cells showed some residual MDA5 oligomerization and antiviral cytokine/ISG responses however indicates the involvement of other mechanisms in regulating MDA5 activation. In particular, the K63-linked polyubiquitination of MDA5 in the helicase domain by TRIM65 has been shown to facilitate MDA5 oligomerization and its downstream antiviral signaling (19). Indeed, silencing of endogenous TRIM65 in WT cells led to a reduction in MDA5 oligomerization to the levels of oligomerization observed for *MDA5^K23R/K43R^* knock-in cells, whereas TRIM65 depletion in the *MDA5^K23R/K43R^* knock-in background near-abolished MDA5 oligomerization (***SI Appendix*, Fig. S5*B***). These data suggest that MDA5 CARD ISGylation and helicase K63-linked ubiquitination play synergistic roles in facilitating MDA5 oligomerization, leading to optimal MDA5 activation. Given the role of the helicase domain in the initial binding to dsRNA ligands, it is tempting to speculate that the TRIM65-mediated ubiquitination of MDA5 occurs first and primes oligomerization, while CARD ISGylation amplifies the magnitude of MDA5 oligomeric assembly and downstream signal transduction. However, additional studies are needed to define the temporal aspects and respective roles of the CARD and helicase PTM-events in the MDA5 oligomerization process, and their relationships to other cofactors needed for MDA5 higher-order assembly formation.

A previous study reported that MDA5 undergoes SUMOylation in the CARDs at K43 (23). However, we observed similar levels of MDA5 SUMOylation (and also K63-linked polyubiquitination) in *MDA5^K23R/K43R^* and WT cells. These results indicate that the two lysine residues are specific for ISGylation, although it is possible that a temporal switch of these two PTMs at K43 can occur for fine-tuning the activation state of MDA5. Future studies are warranted to illustrate the dynamics and relative contributions of MDA5 PTMs in physiological (cell-based or *in vivo*) conditions using similar approaches as described herein for MDA5 CARD ISGylation.

Our identification of ISGylation as a physiologically important PTM governing MDA5-mediated immunity highlights its potential for translational applications. Recent studies have demonstrated that MDA5 plays a determining role in the immunogenicity of COVID-19 vaccines, particularly in stimulating humoral and cell-mediated adaptive immune responses (38, 39). Although the involvement of specific PTMs in MDA5 activation by COVID-19 vaccines remains unknown, we postulate that ISGylation plays a role, and modulating MDA5 ISGylation may provide a strategy to enhance vaccine efficacy. Given that ISG15 conjugation to viral proteins typically inhibits their function, and further, since viruses such as SARS-CoV-2 have evolved tactics to actively remove ISGylation from both host and viral proteins (40–44), boosting ISGylation could offer dual benefits via *1)* fortifying MDA5 (and perhaps other sensor such as cGAS) signaling, and *2)* counteracting viral evasion through de-ISGylation. Along these lines, as sensing of endogenous host RNA ligands by MDA5 and *Mda5/Ifih1* gain-of-function mutations underlie certain autoimmune conditions (45–47), exploring the modulation of MDA5 ISGylation as an immunomodulatory approach to mitigate autoinflammation represents an intriguing area for future research. Overall, our findings unveiling a pivotal role of MDA5 CARD ISGylation in effective innate immunity may hold promise for translational application in antiviral design, vaccinology, and autoimmunity.

## Materials and Methods

### Generation of *MDA5^K23R/K43R^* mice

The *Mda5/Ifih1* transgenic mice were generated by introducing the K23R and K43R mutations into the native *Mda5/Ifih1* genomic DNA (*Ifih1*) locus by replacing the WT exon1 with a double mutant exon1 directly in mice using CRISPR-Cas9 and a targeting vector. sgRNA sequences that directed Cas9 nuclease cutting on either side of a *Mda5/Ifih1* exon1 genomic DNA target fragment were identified by the CRISPR algorithm (http://crispor.tefor.net/) and screened with a sgRNA *in vitro* screening system (Clontech). The cut sites for the 5’ sgRNA *Mda5/Ifih1* 1162/rev (CATCGTGAGGTCTCAGGAAA) and the 3’ sgRNA *Mda5/Ifih1* 1652/fw (CGGGTAGGTGTCAATGTAGT) were then used to design a targeting vector containing a 1 kb 5’ arm of homology, a unique *Asc* I site at the cut site of 1162/rev, a double mutant *Mda5/Ifih1* exon1 sequence, a unique *Pme* I site at the cut site of 1652/fw, and a 1 kb 3’ arm of homology. The insertion of the unique sites prevents cutting the targeting vector by Cas9 nuclease. Mixtures of Cas9 nuclease, both sgRNAs and supercoiled targeting vector were microinjected into the pronucleus of C57BL/6J fertilized oocytes by the Case Transgenic and Targeting Facility (Cleveland, OH). Injected fertilized oocytes were transferred to the oviducts of CD1 pseudo-pregnant recipients and the resulting pups were transferred to our laboratory. In genome editing, because of the two sgRNAs in the mixtures, the DNA repair machinery can also resolve the cuts by consecutive nonhomologous end joining, leading to the deletion of the intertwining WT *Mda5/Ifih1* exon1 sequence and resulting in a putative null allele. Animals were therefore screened for both knock-in (KI) and knock-out (KO) genotypes, with the latter serving as the matched control. The *MDA5^K23R/K43R^* and *MDA5^−/−^* founder mice that harbored the transgenic gene expression were then backcrossed to C57BL6/J WT mice (directly bought from the Jackson Laboratory) to generate homozygous *MDA5^K23R/K43R^* and *MDA5^−/−^* mice in the C57BL6/J background. *MDA5^K23R/K43R^* and *MDA5^−/−^* transgenic mice (founder and up to F7 progeny) were screened and validated by genotyping using a three-set PCR scheme amplifying an exon1-containing fragment. The primer pair A (primers 1 and 2) anneals to the WT exon1 junctions, while the primer pair B (primer 3 and 4) is positioned to anneal at the primer 3’ end to the unique *Asc* I and *Pme* I sites flanking the double mutant exon1. The primer pair C (primers 5 and 6) is located in the distal intronic region flanking both WT and double mutant exon1 (see ***Table 1*** for specific primers). Mice were bred and maintained at the Animal Resources Center of the Cleveland Clinic Florida Research and Innovation Center. No growth or behavioral defects were observed for the *MDA5^K23R/K43R^* and *MDA5^−/−^* mice. All mice were housed in a pathogen-free barrier facility with a 12 h dark and light cycle and ad libitum access to a standard chow diet and water. All mice used in this study were not involved in any other experimental procedure study and were in good health status.

**Table 1.**
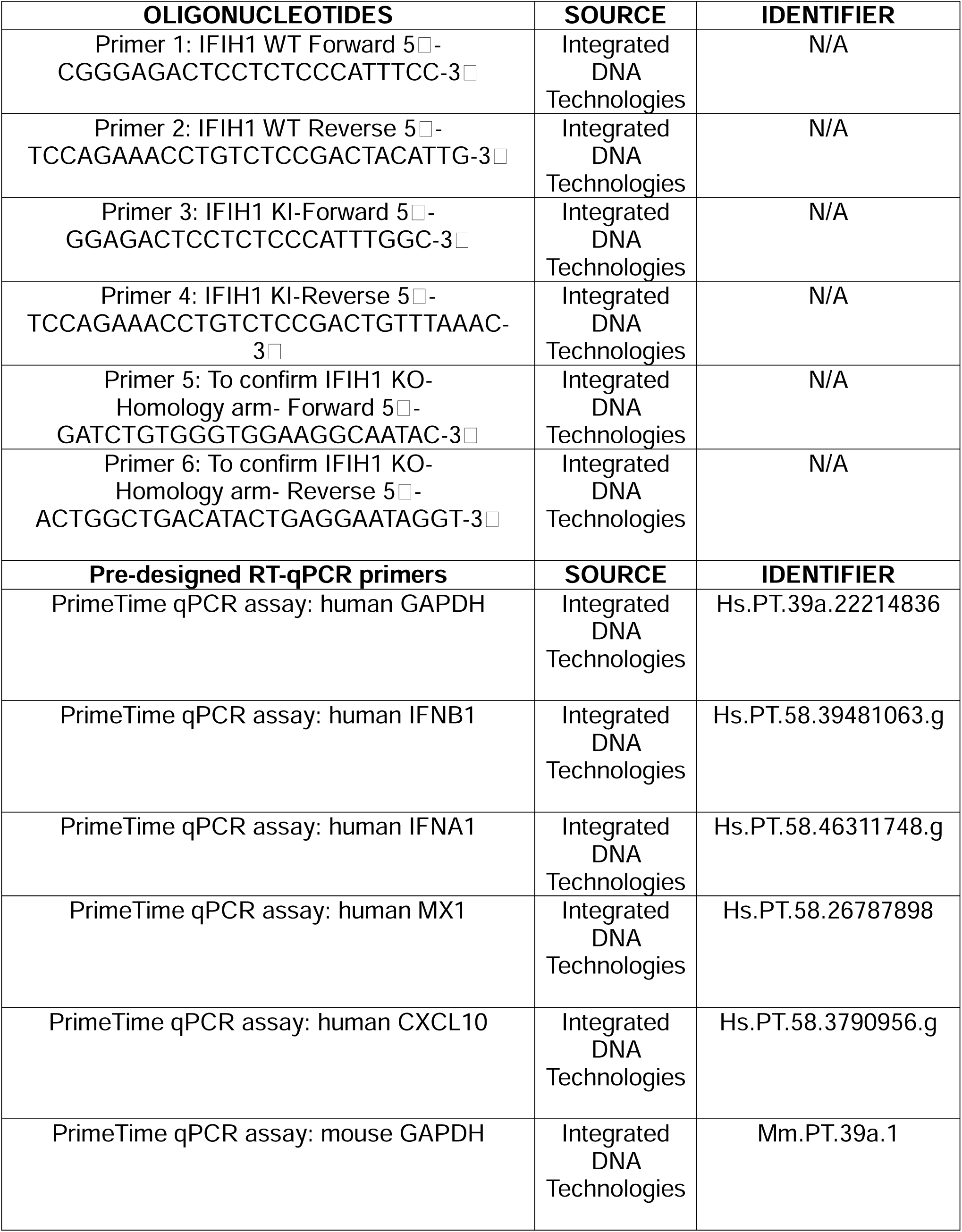
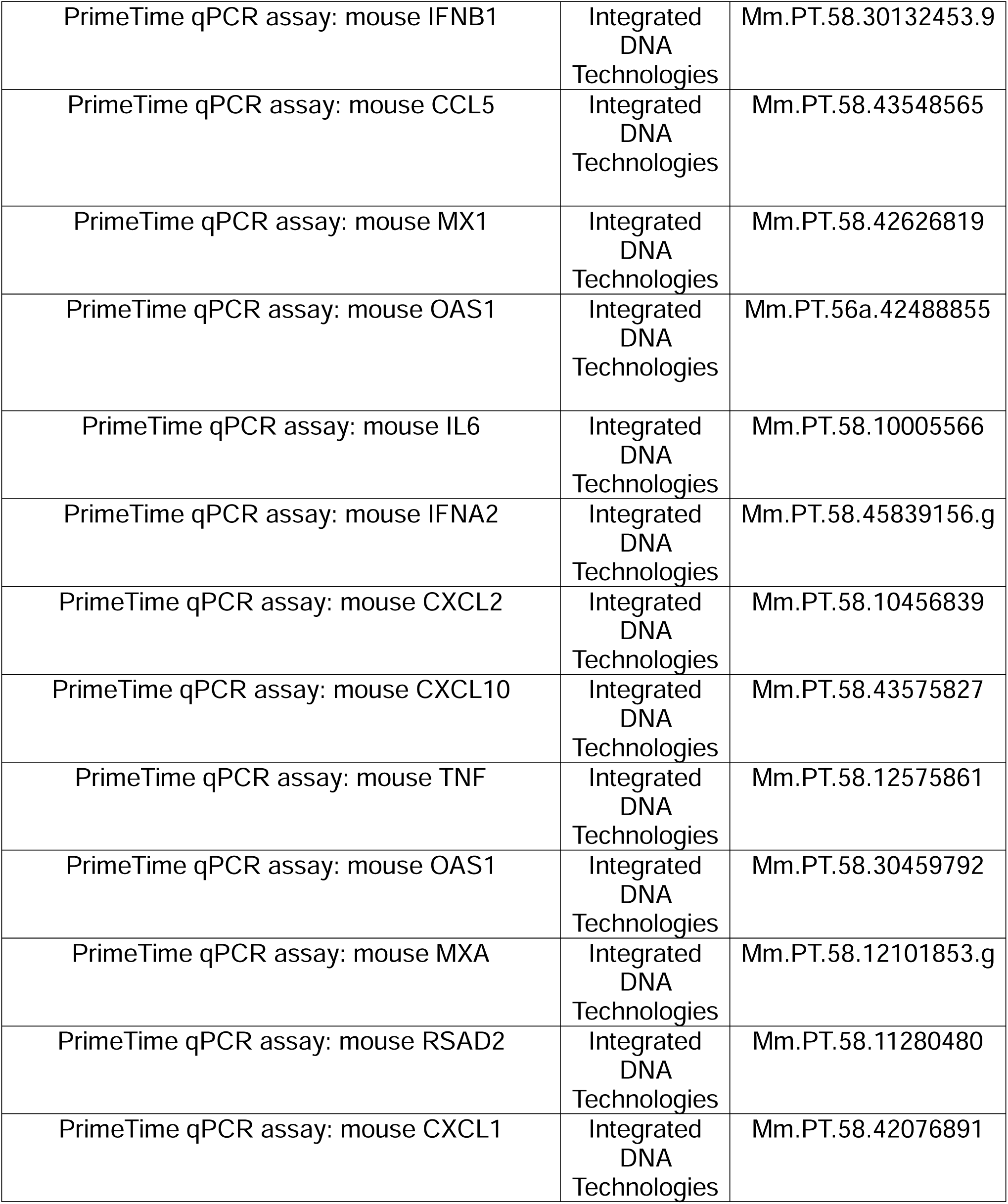

### Mouse infection studies

For EMCV infection, sex-matched, 6-8 week-old WT, *MDA5^K23R/K43R^*, and *MDA5^−/−^* C57BL/6J mice were infected with the indicated plaque forming unit (PFU) of EMCV in 100 µL of sterile PBS via the intraperitoneal route (22, 48–53). Both female and male mice were used in the studies. For survival studies, mice were monitored daily for disease progression, daily signs and symptoms (hind limb paralysis, partial body paralysis, ruffled fur, hunchback, listlessness, trembling, and impaired movement) and euthanized at the indicated times post-infection following humane endpoint criteria defined by Institutional Animal Care and Use Committee guidelines. Retro-orbital blood collection was performed as described previously (54). The blood was centrifuged at 9000 ×*g* for 5 min and stored at −80°C. Whole mouse heart and brain tissues were harvested, longitudinally bisected into two halves, and one half was placed into sterile 1 × PBS, and the other half into TRIzol reagent for RNA isolation and kept on ice. Tissues were homogenized using Qiagen TissueRuptor (22573; Qiagen) at maximum speed for 15 s/sample. Homogenates were clarified by centrifugation at 13,000 ×*g* for 10 min at 4 °C, and supernatants were collected into new sterile tubes and stored at −80 °C (53, 55). EMCV replication in blood, heart, and brain tissues was determined by standard plaque assay (17, 56) or by RT-qPCR analysis of EMCV RNA-dependent RNA polymerase (RdRp; 3Dpol) transcripts using forward primer sequence 5 - GTCATACTATCGTCCAGGGACTCTAT-3 and reverse primer sequence 5 - CATCTGTACTCCACACTCTCGAATG-3 (57). All experiments were performed under protocols approved by the Institutional Animal Care and Use Committee of the Cleveland Clinic Florida Research and Innovation Center.

### Cell culture

HEK293T (human embryonic kidney), primary normal human lung fibroblasts (NHLF), Vero (African green monkey kidney epithelial), and BHK-21 (baby hamster kidney) were purchased from American Type Culture Collection (ATCC) and cultured in Dulbecco’s modified Eagle media (DMEM, Gibco) supplemented with 10% (v:v) fetal bovine serum (FBS, Gibco), 100 U/mL penicillin-streptomycin (Pen-Strep, Gibco), 1 mM sodium pyruvate (Gibco), and 2 mM L-glutamine (Gibco). Vero E6-TMPRSS2 cells were cultured in DMEM supplemented with 10% (v/v) FBS, 1 mM sodium pyruvate, 100 U/mL of penicillin-streptomycin, and 40 µg/mL blasticidin (ant-bl-05; Invivogen). Adult mouse dermal fibroblasts (MDFs) derived from ear/tail tissue of WT*, MDA5^K23R/K43R^*, and *MDA5^−/−^* mice (C57BL/6J mice, 6-8 week-old) were isolated after mincing and then treatment with digestion media containing Collagenase D (20 mg/mL) and Pronase (20 mg/mL) (58, 59). Cells were cultured in DMEM supplemented with 10% (v:v) FBS, 2 mM L-glutamine, 1% (v:v) (NEAA), 1 mM sodium pyruvate, 50 µM 2-mercaptoethanol, and 100 U/ml antibiotic-antimycotic (Gibco). Bone marrow-derived macrophages (BMDMs) were generated from the femur and tibia of WT*, MDA5^K23R/K43R^*, and *MDA5^−/−^* mice (C57BL/6J background, 6-8 week old) and maintained in Roswell Park Memorial Institute (RPMI) media supplemented with 10% (v:v) FBS, 100 U/mL antibiotic-antimycotic (Gibco), 1% (v:v) non-essential amino acids (NEAA), 1 mM sodium pyruvate, and 25 µg/mL macrophage colony-stimulating factor (M-CSF) as previously described (17, 60). All cell cultures were maintained at 37°C in a humidified 5% CO_2_ atmosphere.

Commercially obtained cell lines were authenticated by the respective vendors and were not validated further in the Gack laboratory. Primary WT, *MDA5^K23R/K43R,^* and *MDA5^−/−^* cells were validated by genotyping. Additionally, the presence or absence of MDA5 protein expression was confirmed by IB. All cell lines have been regularly tested for the absence of mycoplasma contamination by PCR assay and/or using the MycoAlert Kit (LT37-701; Lonza).

### Viruses

EMCV (EMC strain, VR-129B) was purchased from ATCC and propagated in HEK293T cells (14). WNV (strain New York 99, NR-158) was purchased from BEI Resources and propagated in Vero cells (56). SeV (strain Cantell) was purchased from Charles River Laboratories. All viral infections were performed by inoculating cells with the virus inoculum diluted in DMEM containing 2% FBS. After 1–2 h, the virus inoculum was removed and replaced with the complete growth medium (DMEM containing 10% FBS) and cells were further incubated for the indicated times. Viral titers in mouse heart and brain homogenates were determined by plaque assay on BHK-21 cells as described previously (53). The plaques were counted, evaluated as PFU/mL [(plaques/well) x (dilution factor)/ (infection volume)], and finally plotted as PFU per gram of tissue (17, 53). Recombinant SARS-CoV-2 (strain K49), propagated in Vero E6-TMPRSS2 cells, was used to isolate RNA for *in vitro* transfections to stimulate MDA5 activation. The SARS-CoV-2 K49 strain was rescued from a bacterial artificial chromosome encoding hCoV-19/Germany/BY-pBSCoV2-K49/2020 (GISAID EPI_ISL_2732373) (61), which was a kind gift from Armin Ensser (Friedrich-Alexander University Erlangen-Nürnberg, Germany). All work with viruses was conducted under approved protocols in the BSL-2/ABSL-2 or BSL-3 facility at the Cleveland Clinic Florida Research and Innovation Center in accordance with institutional biosafety committee regulations and National Institutes of Health (NIH) guidelines.

### Antibodies and other reagents

Primary antibodies used in the present study include anti-MDA5 (1:1,000, D74E4; CST), anti-RIG-I (1:1,000, D14G6; CST), anti-ISG15 (1:500, F-9; Santa Cruz), anti-IFIT2 (1:500, F-12; Santa Cruz), anti-SUMO-1 (1:500, C9H1; CST), anti-K63-Ub (1:500, D7A11; CST), anti-Phospho-IRF3 (Ser396) (1:1,000, D6O1M; CST), anti-IRF3 (1:1,000, D6I4C; CST), anti-Phospho-STAT1 (Tyr701) (1:1,000, 58D6; CST), anti-STAT1 (1:1,000, 9172; CST), anti-Phospho-TBK1 (pSer172) (1:1,000, D52C2; CST), anti-TBK1 (1:1,000, D1B4; CST), anti-HERC5 (1:1,000, 8H23L10; Invitrogen), anti-TRIM65 (1:1,000, HPA021578; Sigma-Aldrich), anti-TRIM25/EFP (1:1,000, 2/EFP; BD Biosciences), anti-ARIH1 (1:2,000, 14949-1-AP; Proteintech), anti-HERC6 (1:1,000, bs-15463R-HRP; Biossusa), anti-UBE1L (1:1,000, JE50-55; Invitrogen), anti-UB2E2 (1:1,000, NBP1-92556; Novus biologicals), anti-Rabbit IgG (1:500, DA1E; CST), and anti-β-actin (1:1,000, C4; Santa Cruz). Anti-mouse and anti-rabbit horseradish peroxidase-conjugated secondary antibodies (1:2,000) were purchased from CST [Anti-mouse IgG, HRP-linked antibody Cell Signaling Technology (#7076), and Anti-rabbit IgG, HRP-linked antibody (#7074)]. Protein G Dynabeads (10003D; Invitrogen) were used for protein IP. Protease (P2714; Sigma Aldrich) and phosphatase inhibitors (P5726; Sigma Aldrich) were obtained from MilliporeSigma. Universal Type I IFN (IFN-α) (11200, PBL Science) was used to stimulate WT, *MDA5^K23R/K43R^*, and *MDA5^−/−^* MDF cells.

### Enzyme-linked immunosorbent assay (ELISA)

For *in vitro* studies, mouse IFN-β protein in the culture supernatants of MDFs from WT, *MDA5^K23R/K43R^*, and *MDA5^−/−^* mice was determined by ELISA using the VeriKine Mouse Interferon Beta ELISA Kit (42400-1; PBL Assay Science) as previously described (14, 17). For *in vivo* studies, mouse IFN-β protein amounts in plasma samples were determined by VeriKine-HS^TM^ Mouse Interferon Beta ELISA Kit (42410-1; PBL Assay Science) following the manufacturer’s instructions (53).

### Viral RNA purification and transfection

EMCV RNA was produced as previously described (20). Briefly, Vero cells were infected with EMCV (MOI 2) for 10 h, and total RNA was isolated using TRIzol Reagent (15596018, Thermo Fisher Scientific) per the manufacturer’s instructions (62, 63). Mock RNA and SARS-CoV-2 RNA were generated by isolating total RNA from Vero E6-TMPRSS2 cells that remained uninfected or that were infected for 24 h with recombinant SARS-CoV-2 (strain K49) (MOI 1) as detailed in previous publications (20, 64). EMCV RNA and SARS-CoV-2 RNA transfections were performed at the indicated concentrations using the Lipofectamine 2000 transfection reagent (11668019; Thermo Fisher Scientific). RABV_Le_ was generated by *in vitro* transcription using the MEGAshortscript T7 Transcription Kit (Invitrogen) according to a previously described protocol (24), and for its transfection into cells, Lipofectamine RNAiMAX Transfection Reagent (13778150; Invitrogen) was used (see Figure legends for details on RABV_Le_ concentrations used).

### Immunoprecipitation assay and Immunoblot analysis

Immunoprecipitation of endogenous proteins (*i.e.,* MDA5, SUMO1) was performed using previously described protocols with minor modifications (14, 20, 65). For assaying endogenous MDA5 ISGylation in MDFs from WT and *MDA5^K23R/K43R^* mice or in primary NHLFs, cells were stimulated as indicated and then lysed using Nonidet P-40 (NP-40) buffer (50 mM HEPES [pH 7.2-7.5], 200 mM NaCl, 1% (v:v) NP-40, 5 mM EDTA, 1× protease inhibitor), followed by centrifugation at 16,000 ×*g* and 4°C for 20 min. Centrifuged cell lysates were then pre-cleared at 4°C for 1-2 h using Protein G Dynabeads pre-conjugated with rabbit IgG (DA1E; CST). Next, cell lysates were incubated with Protein G Dynabeads pre-conjugated with anti-MDA5 antibody (D74E4; CST), or IgG isotype control, at 4°C for 16 h. The beads were extensively washed five times with NP-40 buffer. The proteins were eluted by heating in 1× Laemmli SDS sample buffer at 95°C for 5 min. Protein samples were resolved on Bis-Tris SDS-polyacrylamide gel electrophoresis (PAGE) gels and transferred onto polyvinylidene difluoride (PVDF) membranes (1620177; Bio-Rad). Protein signals were visualized using the SuperSignal West Pico PLUS or Femto chemiluminescence reagents (both Thermo Fisher Scientific) on an ImageQuant LAS 4000 Chemiluminescent Image Analyzer (General Electric) as previously described (20, 66).

For determining the K63-linked ubiquitination and SUMOylation of endogenous MDA5, cell lysates were prepared in a modified RIPA buffer (50 mM Tris-HCl [pH 7.5], 150 mM NaCl, 1% (v:v), NP-40, 2% (w:v) SDS, 0.25% sodium deoxycholate, 1 mM EDTA) followed by boiling at 95°C for 10 min and sonication. The lysates were then diluted 10-fold with the modified RIPA buffer containing no SDS (final concentration of SDS at 0.2%) and cleared by centrifugation at 20,000 ×*g* for 20 min at 4°C. The lysates were pre-cleared as described above, and then subjected to anti-MDA5 (D74E4; CST) or anti-SUMO-1 antibody (C9H1; CST), or IgG (isotype control), following the same protocol as described above (19, 23, 56).

### Knockdown mediated by siRNA

Transient knockdown in primary MDFs or NHLFs was performed using ON-TARGETplus small interfering (si)RNAs (Horizon Discovery) targeting the respective mouse or human genes. These are murine *Herc6* (L-056204-01-0010), murine *Ube2l6* (L-055578-01-0010), murine *Uba7* (L-040733-01-0010), murine *Trim65* (L-058092-01-0010), human *HERC5* (005174-00-0005), human *TRIM65* (L-018490-00-0005), human *TRIM25* (L-006585-00-0005), and human *ARIH1* (L-019984-00-0005). ON-TARGETplus Non-targeting Control Pool (D-001810-10-20) was used as control. Transfection of siRNAs was performed using the Lipofectamine RNAiMAX Transfection Reagent (13778150; Invitrogen) as per the manufacturer’s instructions (17, 20). The knockdown efficiency of the specific genes was determined by RT-qPCR and/or at the protein level by IB using specific antibodies.

### RT-qPCR

Total RNA was purified from indicated cells using the E.Z.N.A. HP Total RNA Kit (Omega Bio-tek) per the manufacturer’s instructions. The quality and quantity of the extracted RNA were assessed using a NanoDrop Lite spectrophotometer. One-step RT-qPCR was performed using the SuperScript III Platinum One-Step RT-qPCR Kit (Invitrogen) with ROX and predesigned PrimeTime qPCR Probe Assays (Integrated DNA Technologies) on a QuantStudio 6 Pro Real-Time PCR System (Applied Biosystems). The relative mRNA expression of the gene of interest was normalized to the levels of cellular *GAPDH* and expressed relative to the values for control cells using the ΔΔCt method. The RT-qPCR primers are listed in **Table 1**.

### Semi-denaturing detergent agarose gel electrophoresis

Endogenous MDA5 oligomerization in EMCV RNA-stimulated MDFs isolated from WT and *MDA5^K23R/K43R^* mice were determined by semi-denaturing detergent agarose gel electrophoresis (SDD–AGE) as previously described (20).

### Sequence alignments

Primary sequence alignment of the amino acid region containing K23 and K43 in orthologous MDA5 proteins was performed using Clustal Omega (1. 2. 4).

### Quantification and Statistical Analysis

All data were analyzed using GraphPad Prism software (version 10). A two-tailed, unpaired Student’s *t*-test was used to compare differences between the two experimental groups in all cases. For statistical evaluation of mice survival, the Log-Rank (Mantel-Cox) test was performed. For the body weight analysis curve, two-way ANOVA was used followed by Bonferroni’s post-test. Significant differences are denoted by \**P* < 0.05, \*\**P* < 0.01, \*\*\**P* < 0.001, and \*\*\*\**P* <0.0001. Pre-specified effect sizes were not assumed, and the number of independent biological replicates (*n*) is indicated for each dataset.

## Acknowledgments

We thank David LePage and Ron Conlon at the Case Transgenic and Targeting Facility of Case Western Reserve University, Ohio, for support with generating the MDA5 knock-in mice. This work was supported by NIH grant R37 AI087846 (to M.U.G).

---

**Figure S1. Validation of *MDA5^K23R/K43R^* mice, and functional assessment of MDA5 oligomerization in cells derived from these mice. *(A)*** Amino acid sequence alignment of the region that contains K23 and K43 (red) in MDA5 from the indicated species using Clustal Omega (1. 2. 4). Numbers denote amino acids. Asterisks define a single, fully conserved residue. Colons (:) indicate conserved groups having strongly similar properties. ***(B)*** Sanger sequencing chromatograms for the *Mda5/Ifih1* exon1 target site in representative WT and *MDA5^K23R/K43R^* mice. The red rectangles indicate the nucleotides encoding the target residues K23 (AAA) and K43 (AAA) (denoted by grey inverted triangles) in WT mice (upper panel), and the introduced one-nucleotide changes to mutate K23 and K43 to arginines (AGA) in *MDA5^K23R/K43R^* mice (middle panel). *Asc* I and *Pme* I are the two unique cut sites flanking the *Mda5/Ifih1* exon1 genomic DNA target. Lower panel: The deletion of the entire exon 1-containing genomic region due to non-homologous end joining (NHEJ) led to the generation of *MDA5^−/−^* mice. ***(C)*** Oligomerization of endogenous MDA5 in primary MDFs isolated from WT and *MDA5^K23R/K43R^* mice that were transfected *ex vivo* with increasing doses of EMCV RNA (0.2 - 0.6 μg/mL) for 16 h, assessed by SDD-AGE and IB with anti-MDA5. MDA5 protein abundance was determined by SDS-PAGE and IB with anti-MDA5. ***(D)*** Densitometric analysis of the MDA5 oligomer signal, normalized to the respective MDA5 abundance, from the experiment in (C). Values represent relative signal intensity normalized to values for unstimulated WT control cells, set to 1. Data are representative of at least three (C and D) independent experiments (mean ± s.d. of n = 3 biological replicates). **P < 0.01, and ****P < 0.0001 (two-tailed, unpaired student’s *t*-test). NS, statistically not significant. CARD, caspase activation, and recruitment domain; CTD, C-terminal domain. dsRNA, double-stranded RNA. Parts of Figure S1A were created using *Biorender.com*.

**Figure S2. The antiviral signaling ability of mouse MDA5 in primary dermal fibroblasts relies on MDA5 ISGylation. *(A)*** IRF3 and TBK1 phosphorylation in WT and *MDA5^K23R/K43R^* mouse-derived MDFs that were infected for 6 h with EMCV (MOI 2) or SeV (250 HAU/mL), assessed in the WCLs by IB with anti-pS396-IRF3, anti-IRF3, anti-pS172-TBK1, and anti-TBK1. ***(B−D)*** Transcript levels of the indicated antiviral or proinflammatory genes in WT, *MDA5^K23R/K43R^*, and *MDA5^−/−^* mouse-derived MDFs that were transfected with EMCV RNA (0.4 μg/mL) or RABV_Le_ RNA (1 pmol/mL) for the indicated times, assessed by RT-qPCR analysis. ***(E−I)*** Transcript levels of the indicated cytokines or ISGs in WT, *MDA5^K23R/K43R^*, and *MDA5^−/−^* mouse-derived MDFs that were infected with EMCV (MOI 1) or SeV (20 HAU/mL) for the indicated times, determined by qRT-PCR. Data are representative of at least two independent experiments (mean ± s.d. of n = 3 biological replicates in (B−I)). *P < 0.05, **P < 0.01, ***P < 0.001, and ****P < 0.0001 (two-tailed, unpaired student’s *t*-test). Red and blue asterisks in (B−I) indicate the statistical significance (P-values) for WT vs. *MDA5^K23R/K43R^* and WT vs. *MDA5^−/−^* values, respectively. h.p.t., hours post-transfection; h.p.i., hours post-infection.

**Figure S3. MDA5 ISGylation promotes MDA5-mediated innate signaling events in immune cells. *(A)*** STAT1 phosphorylation in WT, *MDA5^K23R/K43R^*, and *MDA5^−/−^* mouse-derived BMDMs that were infected with EMCV (MOI 5) or SeV (200 HAU/mL) for 8 h, assessed in the WCLs by IB with anti-pY701-STAT1 and anti-STAT1. ***(B−C)*** *Ifna1* and *Ccl5* mRNA expression in WT, *MDA5^K23R/K43R^*, and *MDA5^−/−^* mouse-derived BMDM*s* that were transfected with EMCV RNA (0.4 μg/mL) or RABV_Le_ RNA (1 pmol/mL) for the indicated times, assessed by RT-qPCR. Data are representative of at least two independent experiments (mean ± s.d. of n = 3 biological replicates in (B−C)). **P < 0.01, ***P < 0.001, and ****P < 0.0001 (two-tailed, unpaired student’s *t*-test). Red and blue asterisks in (B−C) indicate the statistical significance (P-values) for WT vs. *MDA5^K23R/K43R^* and WT vs. *MDA5^−/−^* values, respectively. h.p.t., hours post-transfection.

**Figure S4. HERC5/HERC6-induced MDA5 ISGylation promotes antiviral transcript expression. *(A)*** Silencing efficiency of endogenous *HERC5* and *ARIH1* in primary NHLFs that were transfected for 48 h with the indicated siRNAs and then either Mock-treated or transfected with EMCV RNA (0.4 μg/mL) for 16 h, assessed by RT-qPCR analysis. ***(B)*** *Ifnb1, Ifna1, Rsda2,* and *Tnf* transcripts in WT mouse-derived MDFs that were transfected for 48 h with the indicated siRNAs and then either Mock-treated or transfected with EMCV RNA (0.4 μg/mL) for 16 h, determined by RT-qPCR. The silencing efficiency of endogenous *Herc6* was also evaluated by RT-qPCR analysis. Data are representative of at least two independent experiments (mean ± s.d. of n = 3 biological replicates (A−B)). **P < 0.01, ***P < 0.001, and ****P < 0.0001 (two-tailed, unpaired student’s *t*-test).

**Figure S5. Increased weight loss of *MDA5^K23R/K43R^* mice after EMCV infection as compared to WT mice, and synergistic role of MDA5 regulation by TRIM65 and CARD ISGylation in promoting MDA5 higher-order assemblies.** *(A)* WT, *MDA5^K23R/K43R^*, and *MDA5*^−/−^ mice (6-8-week-old) were infected with EMCV (25 PFU) via *i.p.* inoculation. Body weights of mice were analyzed at the indicated times. *(B)* Endogenous MDA5 oligomerization in WT and *MDA5^K23R/K43R^* mouse-derived MDFs that were transfected for 48 h with either si.C or TRIM65-specific siRNA (si.TRIM65) and then transfected with EMCV RNA (0.4 μg/mL) for 16 h, assessed by SDD-AGE and IB with anti-MDA5. MDA5 protein abundance as well as knockdown of endogenous TRIM65 were determined by SDS-PAGE and IB with anti-MDA5 or anti-TRIM65. Data are representative of at least two independent experiments (mean ± s.d. of n = 6 biological replicates (A)). *P < 0.05, ****P < 0.0001 (Two-way ANOVA followed by Bonferroni’s post-test).

