## Supplementary figures and images for "MDA5 ISGylation is crucial for immune signaling to control viral replication and pathogenesis"

### Supplementary Figure 1

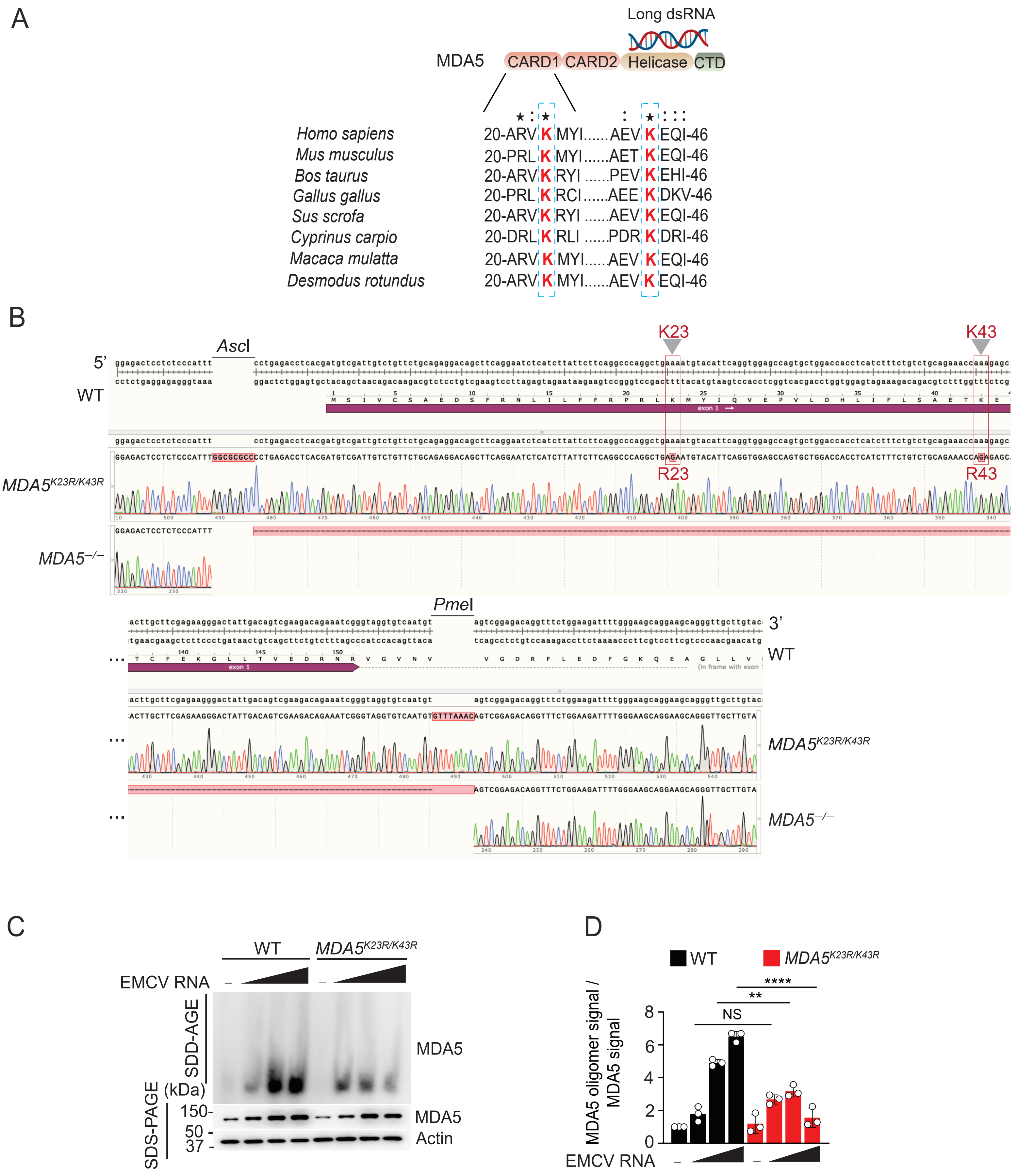

### Supplementary Figure 2

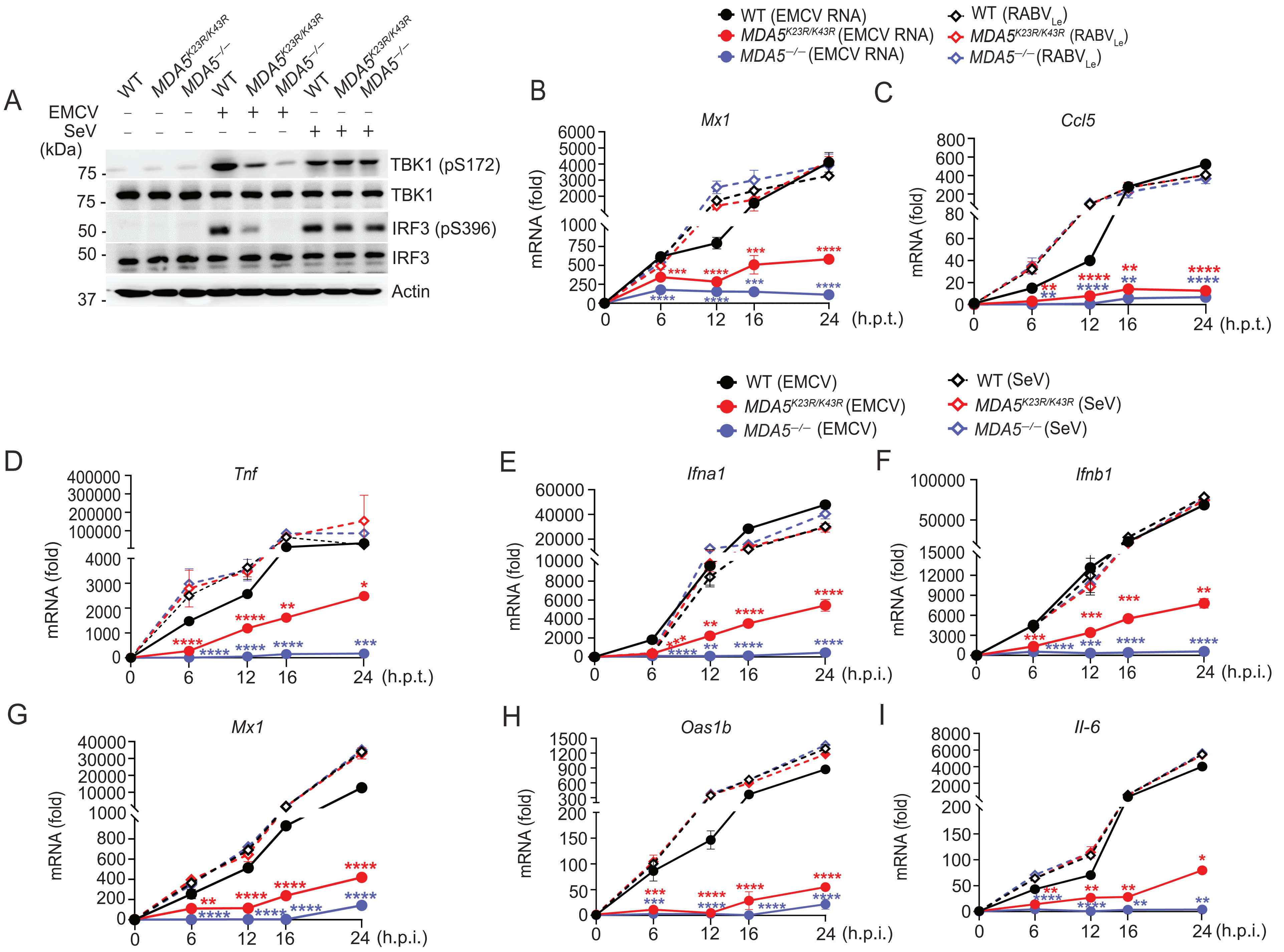

### Supplementary Figure 3

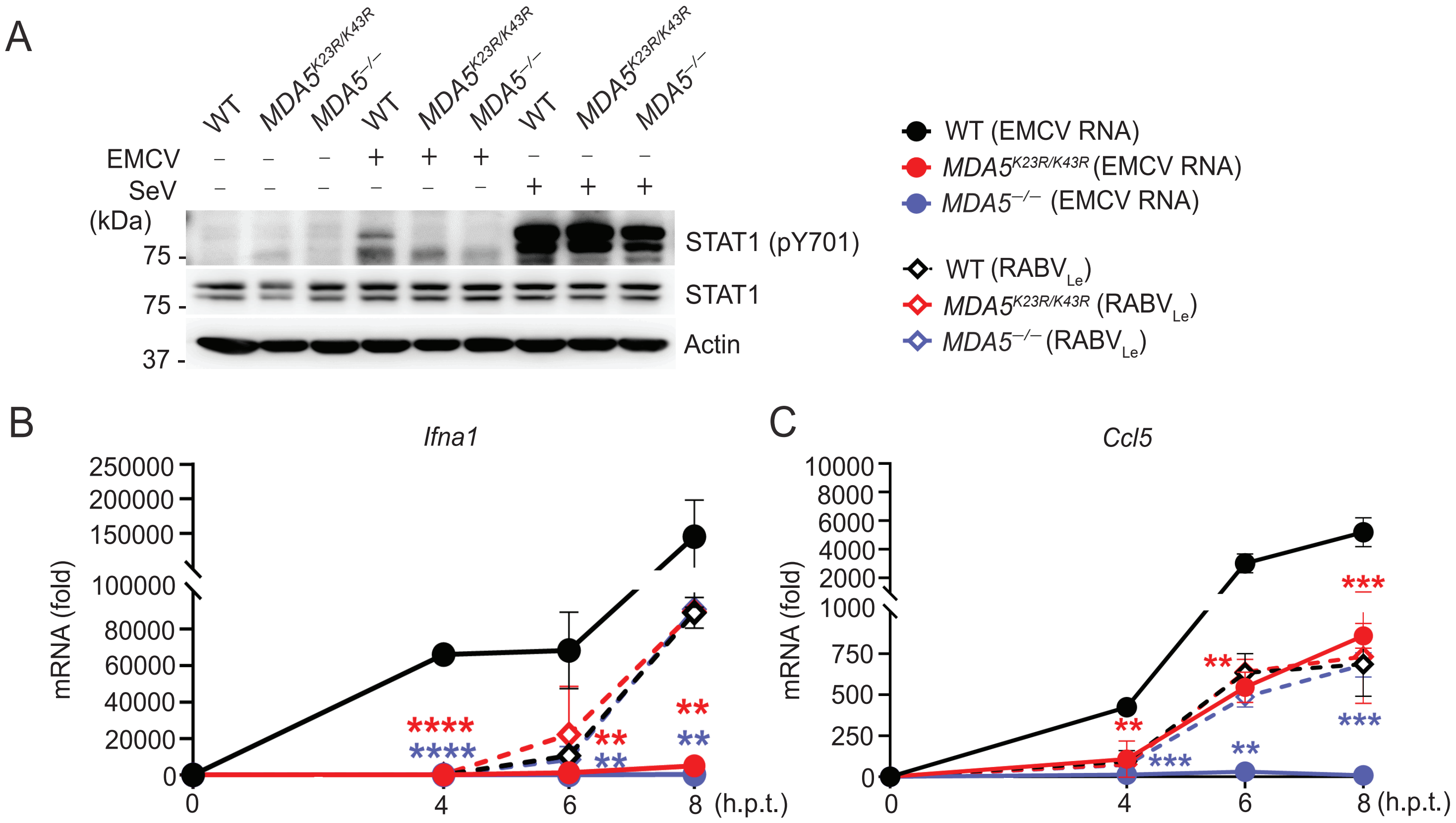

### Supplementary Figure 4

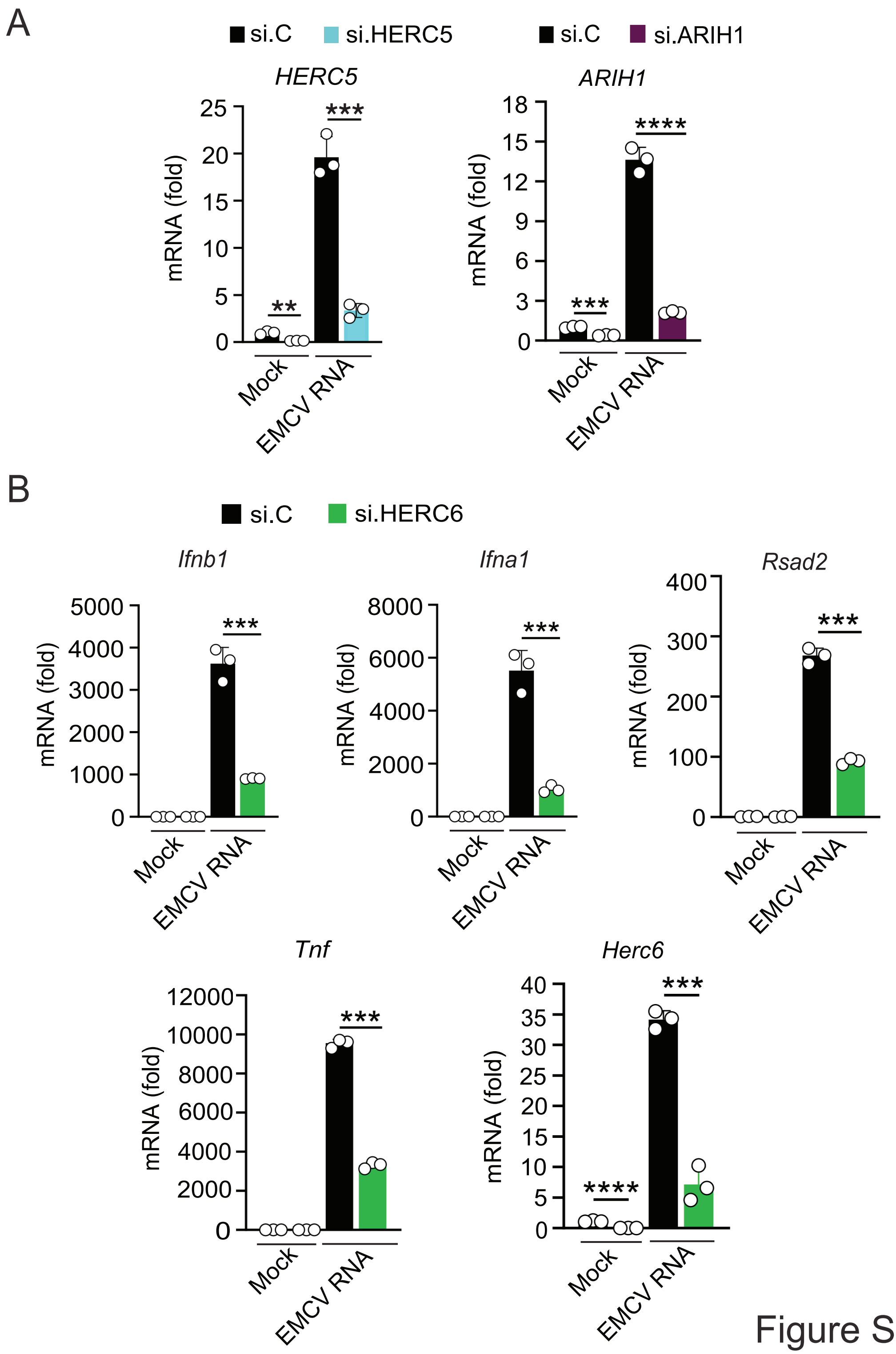

### Supplementary Figure 5

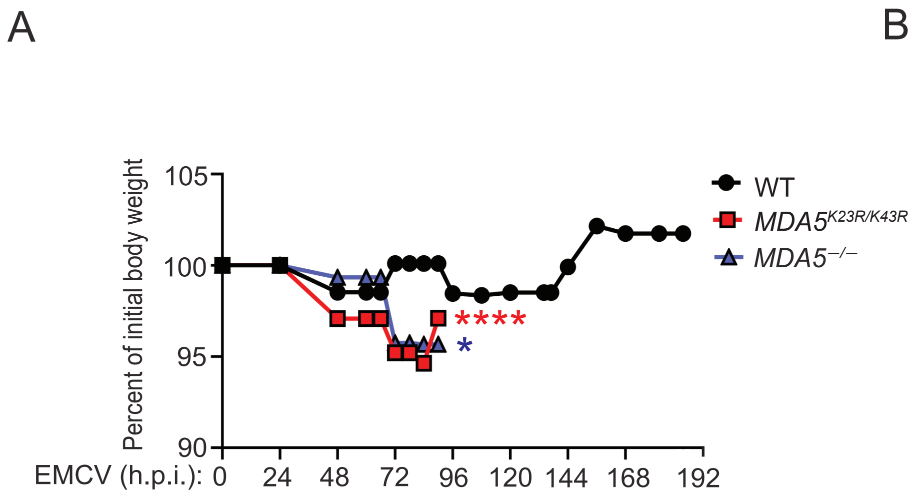

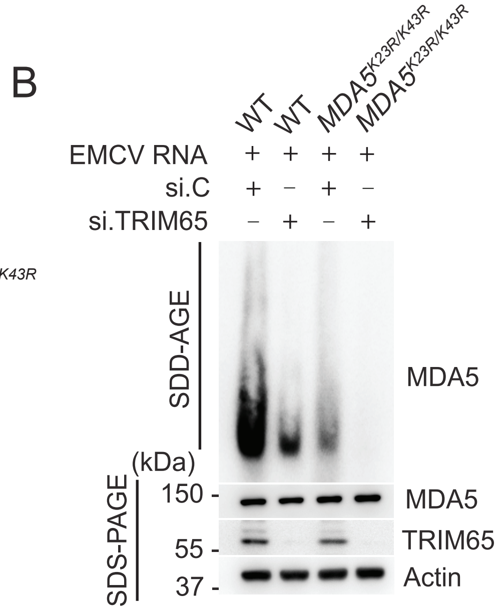
